# The EF-hand domain of MINDY3 is a Ubiquitin and RAD23 UBL-binding domain

**DOI:** 10.1101/2025.07.16.665128

**Authors:** Lee A. Armstrong, Matthew R. McFarland, Rachel O’Dea, Roscislaw Krutyholowa, Magdalena Gorka, Thomas Carroll, Sebastian Glatt, Yogesh Kulathu

## Abstract

The MINDY family of deubiquitinases (DUBs) are exemplified by their preference for cleaving K48-linked polyubiquitin. MINDY3 is architecturally distinct from other MINDY DUBs as its catalytic domain spans the entire length of the protein except for an atypical EF-hand insertion. We uncover this EF-hand (MINDY3^EF-hand^) to be a ubiquitin-binding domain with three distinct binding sites, enabling MINDY3 to bind and effectively cleave long polyubiquitin chains. Furthermore, the MINDY3^EF-hand^ domain binds not only to polyubiquitin but also to the UBL domain of the proteasome shuttling and DNA repair factors RAD23A and RAD23B. The MINDY3^EF-hand^ facilitates this interaction with RAD23s in cells and mediates MINDY3 recruitment to DNA damage sites, establishing this unique DUB as a potential regulator of cellular DNA damage responses. MINDY3 binds specifically to the UBL domain of RAD23s, and none of the other UBLs tested. The crystal structure of the MINDY3^EF-hand^:RAD23A^UBL^ domain complex reveals the molecular basis for specificity. We find that MINDY3 can form a ternary complex with RAD23A/B and polyubiquitin, and our findings suggest a model wherein MINDY3 can deubiquitylate RAD23A/B-bound clients.

## Introduction

Ubiquitylation is a posttranslational modification (PTM) involved in all aspects of eukaryotic cell biology. Canonical ubiquitylation occurs through the formation of an isopeptide bond between the C-terminus of a ubiquitin molecule and a surface lysine residue on the target protein. This conjugated ubiquitin monomer can then be extended by the conjugation of additional ubiquitin monomers onto one of its seven lysine residues (K6, K11, K27, K29, K33, K48, K63) or the N-terminal methionine, to form polyubiquitin (polyUb) chains (Yau & Rape, 2016).

These diverse ubiquitin architectures used in cell signalling are regulated by deubiquitinases (DUBs) which can edit ubiquitin chains by trimming them or removing them entirely from substrates. Of the 100 known DUBs in humans, many are promiscuous in their cleavage activities and will cleave polyUb without regard for linkage type. However, some DUB families, such as OTU, JAMM and MINDY, are highly specific at cleaving particular linkage types (Lange *et al*, 2022; Clague *et al*, 2019). This behaviour is dictated by the nature of the interaction between the respective DUB and polyUb. For instance, DUBs like OTUD7B and AMSH require simultaneous binding at both S1 and S1’ ubiquitin binding sites (Mevissen *et al*, 2016; Sato *et al*, 2008). This dual-site binding requirement positions the scissile bond precisely across the catalytic site, resulting in high specificity for cleaving certain linkage types, and simultaneously excluding others.

In addition to binding polyUb with their catalytic core domains, many DUBs possess auxiliary domains that modulate their cellular functions. These domains can play important roles in modulating DUB function, influencing their subcellular localisation as well as interactions with substrates and regulators (Sowa *et al*, 2009; Lange *et al*, 2022). Other auxiliary domains serve as additional Ubiquitin Binding Domains (UBDs), further enhancing or restricting the substrate specificity of the enzyme. For example, OTUD1 lacking its K63-specific UIM domain loses its specificity and exhibits reduced cleavage efficiency (Mevissen *et al*, 2013).

We previously reported the discovery of the MINDY family of DUBs (Abdul Rehman *et al*, 2016), and more recently characterised the mechanisms regulating the activation and activity of MINDY1 and MINDY2 (Abdul Rehman *et al*, 2021). Here, we focus on MINDY3, which shares only 20% sequence similarity with the catalytic domains of MINDY1 or MINDY2, and whose domain architecture is distinct from the other human MINDY DUBs. By structurally and biochemically characterising the mechanisms of polyUb recognition and catalysis, we discover the EF-hand domain in MINDY3 (MINDY3^EF-hand^) serves as a unique ubiquitin binding domain that enables MINDY3 to bind and cleave long polyUb chains. Furthermore, the EF-hand domain interacts with the UBL domain of the proteasome shuttling/DNA repair factors RAD23A and RAD23B and, intriguingly, facilitates the specific recruitment of MINDY3 to sites of DNA damage.

## Results and Discussion

### MINDY3 cleaves K48 linkages in a chain length-dependent manner

MINDY3 has a domain architecture that is distinct from MINDY1 and MINDY2 (MINDY1/2) and consists largely of a catalytic domain with an insertion of a predicted EF-hand (**Fig 1A).** Like MINDY1/2, MINDY3 is specific towards cleaving K48-linked polyUb (Abdul Rehman *et al*, 2016). To determine whether MINDY3 also shares a preference for recognising and cleaving longer polyUb chains as observed in MINDY1/2, we monitored full length (FL) MINDY3 cleavage of K48-linked chains of increasing length (**Fig 1B**). While only ∼40% of K48-Ub2 is cleaved after 90 minutes, K48-Ub5 is completely cleaved after the same incubation time. Therefore, the cleavage efficiency of MINDY3 increases with increased Ub chain length, which mirrors the chain length sensing of MINDY1/2.

**Figure 1.**
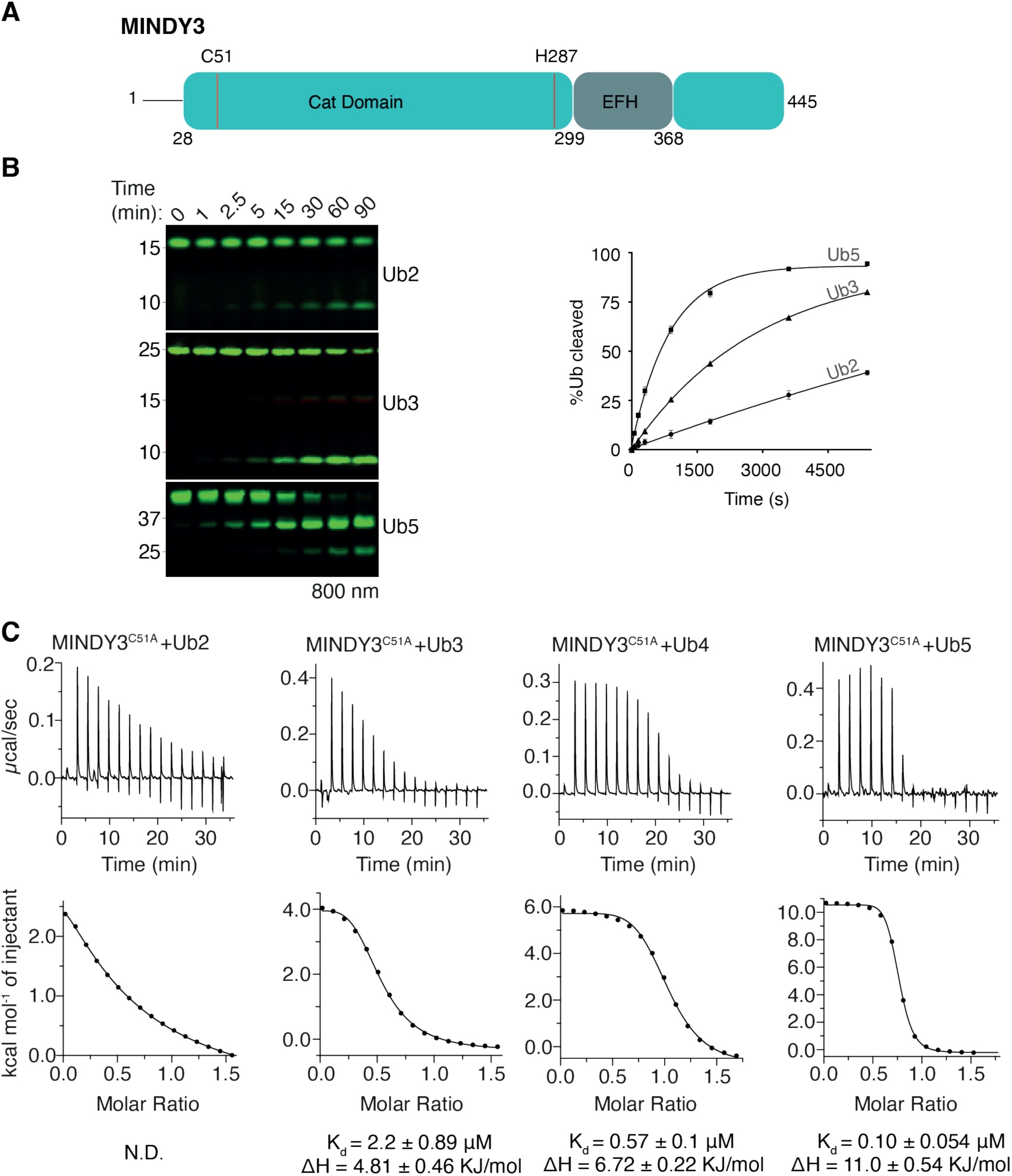
MINDY3 prefers binding to and cleaving long K48-linked polyUb. **(A)** Schematic depicting the domain architecture of MINDY3. **(B)** DUB assay with FL MINDY3 against K48-linked polyUb chains fluorescently labelled at the distal end of the chain. Fluorescence intensity at each time point was measured and plotted to quantify the cleavage of each chain length. **(C)** Isothermal Titration Calorimetry (ITC) measurements in which MINDY3^C51A^ was titrated into K48-linked polyUb chains of varying lengths.

To test how MINDY3 senses ubiquitin chain length, we analysed the binding of catalytically dead MINDY3 (MINDY3^C51A^) to K48-linked chains of varying lengths by size exclusion chromatography (SEC). MINDY3^C51A^ and K48-Ub2 elute as separate species, whilst with K48-Ub3 and K48-Ub4, some DUB-polyUb complex formation can be observed. MINDY3^C51A^ forms a stable complex with K48-Ub5 and the two proteins coelute together (**Fig EV1A**). To quantify the binding parameters, we performed Isothermal Titration Calorimetry (ITC) experiments. We could not observe detectable binding to K48-Ub2, while K48-Ub3, -Ub4 and -Ub5 bind with affinities of 2.2 ± 0.9 μM, 0.57 ± 0.1 μM and 0.1 ± 0.05 μM, respectively (**Fig 1C**). MINDY3’s enhanced binding and cleavage of longer Ub chains suggested the presence of several ubiquitin binding sites within its catalytic site.

To determine how MINDY3 cleaves chains of different lengths, we first analysed the cleavage of K48-linked Ub5. At the earliest time points, the only visible cleavage products are Ub4 and monoUb (Ub1), while shorter products such as Ub2 only appear after longer incubation times (**Fig EV1B**). In contrast, Miy2 (Ypl191c), the yeast homologue of MINDY2 and an endo-DUB, produces cleavage products of all possible chain lengths (**Fig EV1B**). This suggests that MINDY3 works as an exo-DUB, cleaving from one end of the chain, when presented with shorter chains. To analyse cleavage of even longer chains, we incubated MINDY3 with a mixture of K48-linked chain lengths (Ub_6_-Ub_20_). Here, the ladder of polyUb is rapidly cleaved down to predominantly Ub5 with other smaller species present, indicating that chains are cleaved internally in an endo-mode of cleavage. The resultant Ub5 is then subsequently processed by exo-cleavage. Miy2 in contrast cleaves chains of all lengths in an endo-manner (**Fig EV1C**). To conclusively establish this switch in cleavage mode, we incubated MINDY3 with polyUb chains that are fluorescently labelled only on the extreme distal Ub moiety (Abdul Rehman *et al*, 2021). If MINDY3 is a true exo-DUB, the only visible cleavage product should be fluorescent monomer. However, MINDY3 produces a broad-range of smaller fluorescent labelled species indicative of an endo-mode of cleavage (**Fig EV1D**). Hence, MINDY3 like MINDY1/2 cleaves as an exo-DUB from the distal end with shorter polyUb chains (≤ 5 Ubs) and in an endo-fashion with longer polyUb chains (> 5 Ubs) (Abdul Rehman *et al*, 2021). Detecting linkage type and chain length therefore appears to be a shared feature within the MINDY family.

### Crystal structure of MINDY3 reveals an architecture distinct from MINDY1/2

To determine whether a common protein fold between MINDY3 and MINDY1/2 can explain the similarities in catalytic activities and shed light on the role of the EF-hand insertion, we determined the crystal structure of full length MINDY3 (**Fig. 2A**) (**Table 1**). While discernible electron density was present for most of the molecule, electron density for the EF-hand domain (302-376) and a region within the catalytic domain (111-138) was not visible, indicating intrinsic flexibility. While the catalytic domains of MINDY1 and MINDY2 are very similar (Abdul Rehman *et al*, 2021), MINDY3^apo^ shows several differences in comparison to MINDY1(RMSD of 14.9Å _158 Cα_) and MINDY2 (RMSD of 17.3Å _116 Cα_) (**Fig. EV2B**). Indeed, a DALI search for structurally related proteins identifies MINDY4 as the closest match (Z-score 47.2 (MINDY4), 4.2 (MINDY1) (Holm *et al*, 2023)).

**Figure 2.**
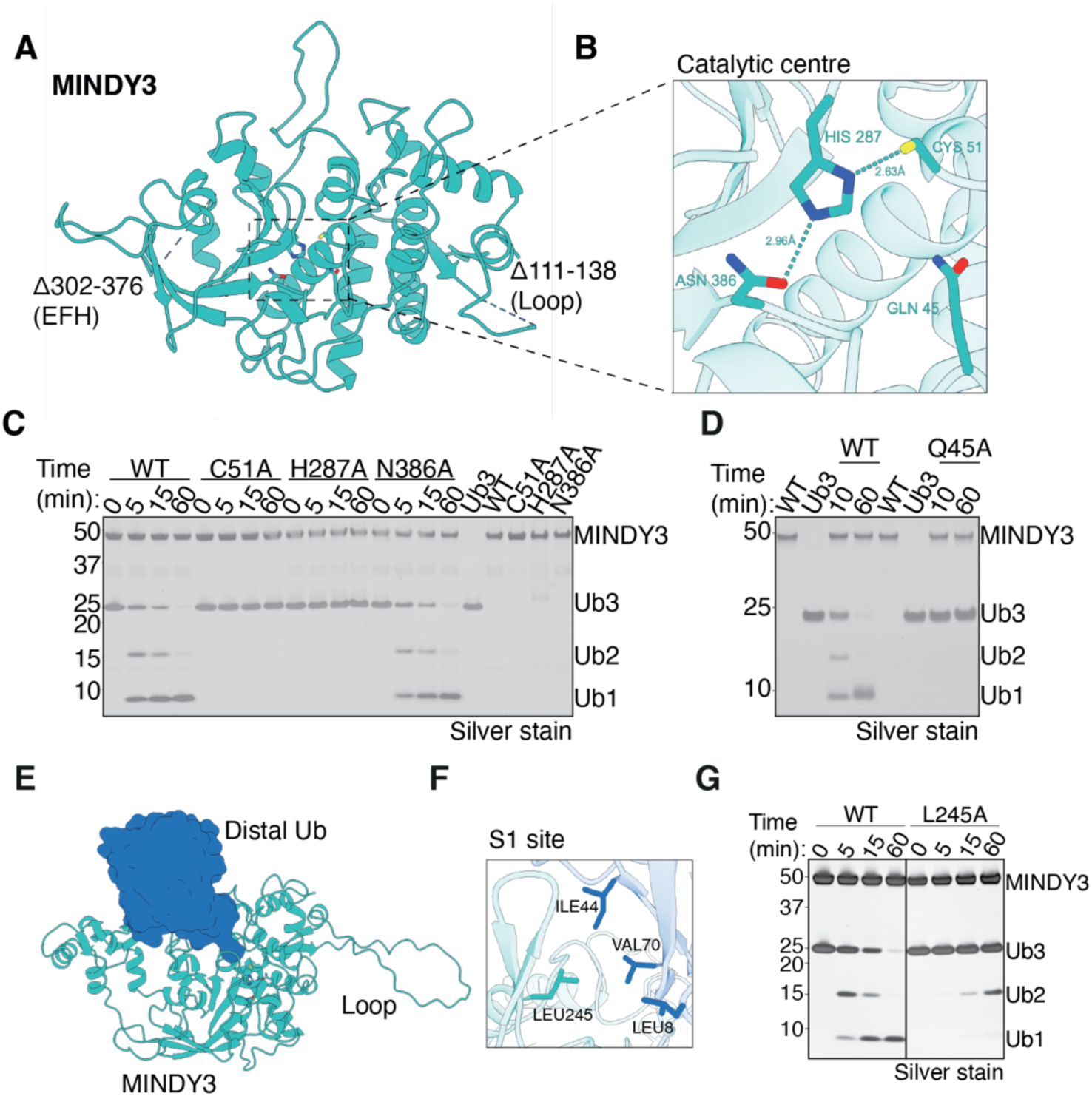
Crystal structure of MINDY3 catalytic domain. **(A)** Cartoon representation of the crystal structure of FL MINDY3. Electron density for the EF-hand (302-376) and a flexible loop within the catalytic domain (111-138) are missing and are indicated by a dotted line on the structure. **(B)** Close up view of the catalytic centre of MINDY3 showing an apparent canonical catalytic triad consisting of C51, H287 and N386. Q45 is positioned to act as the oxyanion hole. DUB assay testing cleavage of K48-Ub3 and indicated alanine mutants of the catalytic triad residues **(C)** and oxyanion hole **(D)**. **(E)** Alphafold model of FL MINDY3 in complex with monoUb. **(F)** Inset shows interaction between MINDY3 Leu245 and the Ile44 patch of ubiquitin. **(G)** DUB assay of MINDY3 and MINDY3^L245A^ against K48-Ub3 chains.

**Table 1.**
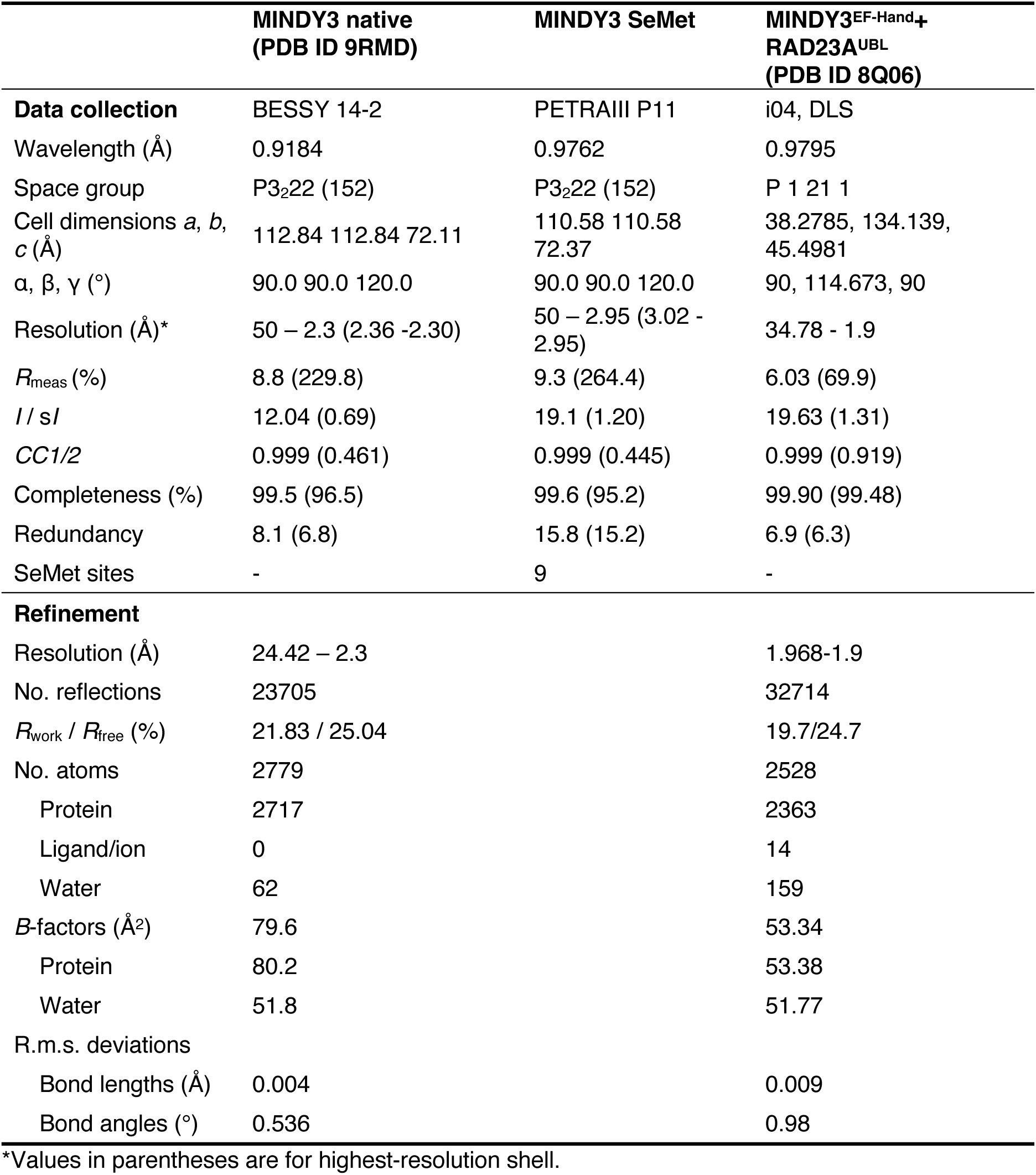
Data collection and refinement statistics.

|  | MINDY3 native<br>(PDB ID 9RMD) | MINDY3 SeMet | MINDY3<sup>EF-Hand+</sup><br>RAD23A<sup>UBL</sup><br>(PDB ID 8Q06) |
| --- | --- | --- | --- |
| **Data collection** | BESSY 14-2 | PETRAIII P11 | i04, DLS |
| Wavelength (Å) | 0.9184 | 0.9762 | 0.9795 |
| Space group | P3<sub>2</sub>22 (152) | P3<sub>2</sub>22 (152) | P 1 21 1 |
| Cell dimensions *a*, *b*,<br>*c* (Å) | 112.84 112.84 72.11 | 110.58 110.58<br>72.37 | 38.2785, 134.139,<br>45.4981 |
| $\alpha$, $\beta$, $\gamma$ (°) | 90.0 90.0 120.0 | 90.0 90.0 120.0 | 90, 114.673, 90 |
| Resolution (Å)* | 50 – 2.3 (2.36 -2.30) | 50 – 2.95 (3.02 -<br>2.95) | 34.78 - 1.9 |
| *R*<sub>meas</sub> (%) | 8.8 (229.8) | 9.3 (264.4) | 6.03 (69.9) |
| *I* / *sI* | 12.04 (0.69) | 19.1 (1.20) | 19.63 (1.31) |
| *CC*1/2 | 0.999 (0.461) | 0.999 (0.445) | 0.999 (0.919) |
| Completeness (%) | 99.5 (96.5) | 99.6 (95.2) | 99.90 (99.48) |
| Redundancy | 8.1 (6.8) | 15.8 (15.2) | 6.9 (6.3) |
| SeMet sites | - | 9 | - |
| **Refinement** |  |  |  |
| Resolution (Å) | 24.42 – 2.3 |  | 1.968-1.9 |
| No. reflections | 23705 |  | 32714 |
| *R*<sub>work</sub> / *R*<sub>free</sub> (%) | 21.83 / 25.04 |  | 19.7/24.7 |
| No. atoms | 2779 |  | 2528 |
| Protein | 2717 |  | 2363 |
| Ligand/ion | 0 |  | 14 |
| Water | 62 |  | 159 |
| *B*-factors (Å<sup>2</sup>) | 79.6 |  | 53.34 |
| Protein | 80.2 |  | 53.38 |
| Water | 51.8 |  | 51.77 |
| R.m.s. deviations |  |  |  |
| Bond lengths (Å) | 0.004 |  | 0.009 |
| Bond angles (°) | 0.536 |  | 0.98 |

\*Values in parentheses are for highest-resolution shell.

In MINDY3, the catalytic cysteine, Cys51 is positioned ∼2.6 Å from the catalytic histidine residue (His287) and is properly aligned for catalysis. A nearby asparagine, Asn386, sits ∼3 Å away from the catalytic histidine and could coordinate His287 performing the function of the negatively charged residue within the catalytic triad (**Fig. 2B**). Alanine substitutions of the catalytic residues, C51A and H287A, completely abolished DUB activity (**Fig. 2C**). Surprisingly however, the N386A mutant remains as active as WT in cleaving Ub3, demonstrating that N386 is dispensable for activity. This suggests that MINDY3, like MINDY1/2 relies on a catalytic dyad for cleavage. Of note, Gln45, which forms the oxyanion hole that stabilises the tetrahedral intermediate, is essential for the DUB activity of MINDY3 (**Fig. 2D**).

We next wanted to understand how ubiquitin engages with MINDY3 at the S1 site. To that end, we generated an AlphaFold model of MINDY3 in complex with monoUb (**Fig. 2E**). The distal Ub sits at the S1 with the C-terminal tail of Ub threaded down into the active site of MINDY3 and is predicted to bind via its Ile44 patch in an interaction centred on Leu245 in MINDY3 (**Fig. 2F**). To test this model, we tested the cleavage of K48-Ub3 by MINDY3^L245A^ mutant in a DUB assay. Indeed, DUB activity was completely abolished (**Fig. 2G**), which highlights both the essentiality of Ub binding at the S1 site and the accuracy of the AlphaFold model.

### MINDY3 has an atypical EF-hand domain

As no discernible electron density was present for the EF-hand in the crystal structure, we predicted the structure of FL MINDY3 using AlphaFold (Jumper *et al*, 2021). The AlphaFold model of the catalytic domain is very similar to the experimentally determined crystal structure (RMSD of 0.484Å _279 Cα_)) (**Fig. 3A, EV2C**). In addition, the AlphaFold model predicts a large, solvent-exposed loop that corresponds to the region missing from the crystal structure (**Fig. EV2D**) and an EF-hand domain that is juxtaposed to the catalytic domain at the end of two flexible linkers. Mapping the evolutionary conservation of each residue onto the AlphaFold model reveals that the catalytic core and EF-hand domain to be highly conserved in vertebrates (Ashkenazy *et al*, 2016) (**Fig. EV2A**). To investigate the contribution of the extended loop and the conserved EF-hand domain to MINDY3 activity, we individually removed these two elements and assessed DUB activity. While removal of the extended loop had no effect on cleavage of K48-Ub3 (**Fig. EV2D**) or longer K48-linked polyUb (**Fig. EV2E**), deletion of the EF-hand domain resulted in a detectable decrease in cleavage activity towards K48-linked Ub3 (**Fig. 3B**), and a substantial decrease in cleavage activity towards longer K48-linked polyUb (**Fig. 3C**).

**Figure 3.**
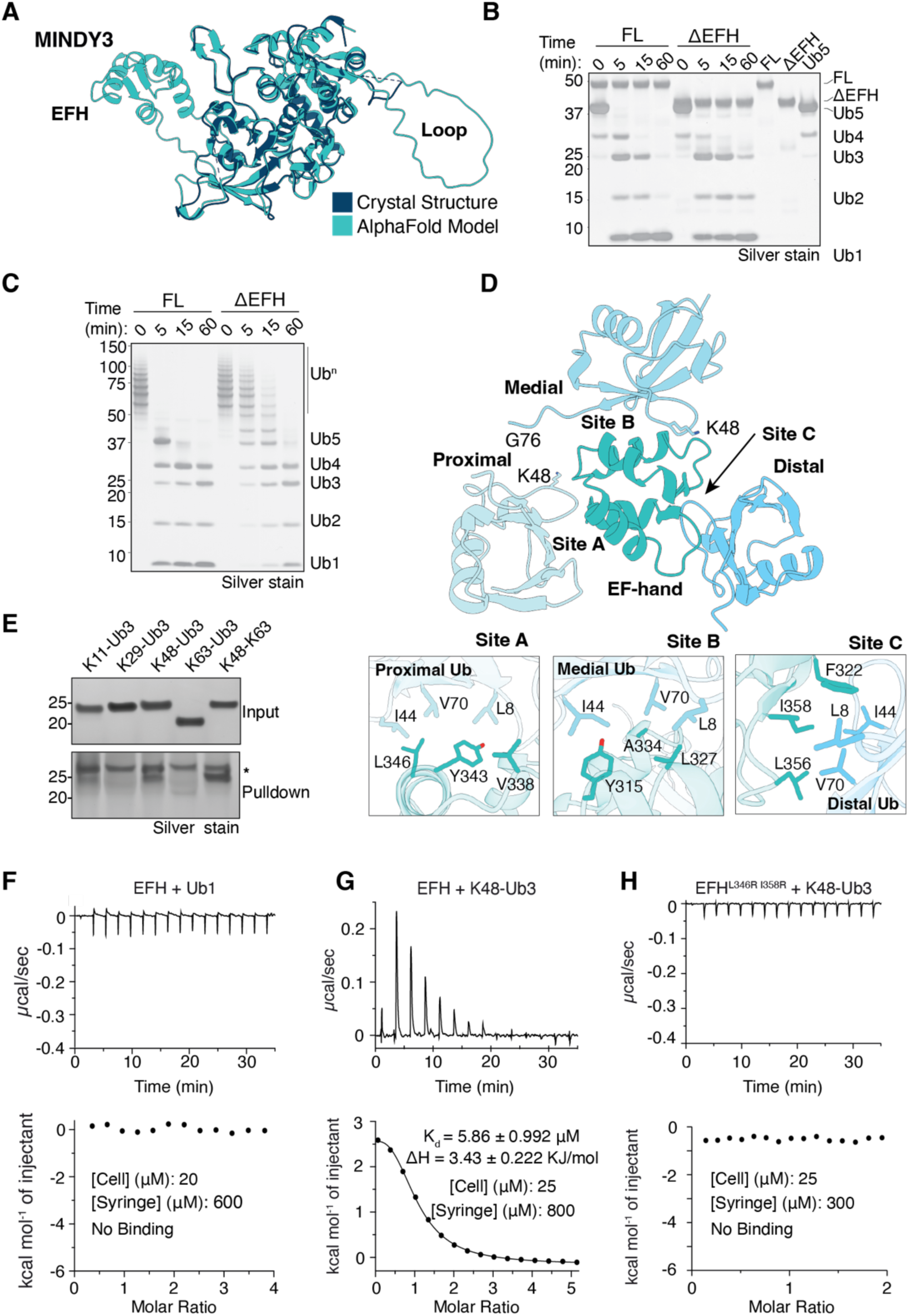
The EF-hand is a ubiquitin binding domain required for cleavage of long polyubiquitin. **(A)** AlphaFold Model of FL MINDY3 superposed onto the experimentally determined crystal structure of MINDY3. The EF-hand domain and the loop not present in the crystal structure are highlighted. DUB assay monitoring cleavage by WT and ΔEF-hand MINDY3 of K48-linked Ub5 **(B)** and long (Ub6-20) K48-linked polyUb **(C)**. **(D)** AlphaFold model of the EF-hand of MINDY3 bound to 3 ubiquitin molecules. The ‘distal’ ubiquitin binds to the EF-hand motif loops. K48 and G76 are highlighted to show how K48-linkages could be formed between the ubiquitin moieties. Inset shows a close-up view of the interactions of the EF-hand with the proximal, middle and distal ubiquitin via Sites A, B and C, respectively. **(E)** Pull down assay where HALO-Tag MINDY3^EF-hand^ protein immobilised on resin was incubated with K11, K29, K48 or K63 homotypic trimers, or a branched K48-K63 trimer. * indicates background band (**F-I**) ITC measurements in which MINDY3^EF-hand^ is titrated into monoUb **(F)** or K48-linked Ub3 **(G)**, or **(I)** an L346R, I358R mutant EF-hand titrated into K48-linked Ub3.

Canonical EF-hand domains are the most common calcium binding domains found in proteins (Holmes, 1996) with roles in several key cellular processes including signal transduction, muscle contraction and calcium homeostasis (Chin & Means, 2000; Schwaller, 2010; Gagné *et al*, 1995; Skelton *et al*, 1994). EF-hand domains typically consist of a helix-loop-helix typically made up of a 12 residue motif: X•Y•Z•(-Y)•(-X)••(-Z) where X, Y, Z and -X, -Y and -Z represent residues that coordinate the metal ion and • represents intervening residues. In most cases, the motifs occur in pairs (Lewit-Bentley & Réty, 2000). Aligning the EF-hand motifs from 878 vertebrate genes gives a consensus sequence for an EF-hand motif (Kuznetsov *et al*, 2023), which reveals that some positions are almost invariant, while others can tolerate certain substitutions (**Fig EV3A**). The MINDY3^EF-hand^ shows significant divergence from the canonical sequence but still harbours invariant residues and preserves the overall biochemical character at specific positions. Most notably, several charged residues, which would be important for metal/calcium coordination are replaced by hydrophobic residues (**Fig. EV3B).** The evolutionary conservation of these non-canonical hydrophobic residues indicates that the inability to bind Ca^2+^ is an evolutionarily conserved trait of MINDY3 (**Fig. EV3C**). As the MINDY3^EF-hand^ domain is required for efficient cleavage of polyUb, we tested whether Ca^2+^ can influence MINDY3 DUB activity after incubating with either CaCl_2_ or EDTA/EGTA. As chain cleavage was unaffected under these conditions (**Fig. EV3D**), it provides further evidence that Ca^2+^ is not required for the function of the MINDY3^EF-hand^ and MINDY3.

### The EF-hand domain of MINDY3 is a UBD

Since the removal of the MINDY3^EF-hand^ domain impacts the cleavage of longer polyUb and several hydrophobic residues are present in Motif 2 of the EF-hand domain, we wondered if the MINDY3^EF-hand^ domain contributes to the cleavage of long polyUb chains by functioning as a UBD. To test this hypothesis, we first used AlphaFold to predict a complex of the MINDY3^EF-hand^ with monoUb. Indeed, the predicted model shows that the MINDY3^EF-hand^ can potentially bind monoUb with an interface formed around the hydrophobic Ile44 patch of Ub (**Fig. 2E**). Next, we performed ITC measurements using purified MINDY3^EF-hand^ and monoUb but could not detect binding (**Fig. 3G**). Analytical SEC experiments also showed two separately eluting species (**Fig. EV3F**). These results suggested that the EF-Hand either does not bind Ub or the affinity is too low for detection by these methods (Hurley *et al*, 2006; Kaiser *et al*, 2011).

We next explored whether the MINDY3^EF-hand^ may have multiple ubiquitin binding sites. Using AlphaFold to predict complexes of the MINDY3^EF-hand^ bound to multiple ubiquitin monomers revealed a striking arrangement where the MINDY3^EF-hand^ domain can bind three ubiquitin molecules simultaneously. In this predicted complex structure, the C-terminus of one Ub points towards K48 of another Ub (**Fig. 3D**). Hence, this arrangement suggests a model where a K48-linked Ub3 chain could bind to the MINDY3^EF-hand^ domain through three distinct binding surfaces, named sites A, B, and C, which would engage the proximal, middle, and distal ubiquitin molecules respectively. The Predicted Aligned Error (PAE) plot for the model shows high confidence for the positioning of the distal and proximal ubiquitin (**Fig. EV3E**). Of note, the Ile44 patches of each Ub is modelled to bind to hydrophobic patches on the EF-hand (**Fig. 3F**).

If this predicted model is correct, then the MINDY3^EF-hand^ should bind to polyUb with higher affinity. Indeed, we could detect complex formation between the isolated MINDY3^EF-hand^ domain and K48-linked Ub3 by SEC (**Fig. EV3G**). Furthermore, ITC measurements reveal the MINDY3^EF-hand^ to bind K48-linked Ub3 with a Kd value of ∼6 μM (**Fig. 3H**). The endothermic enthalpy change indicates that the interaction is driven by favourable entropy, likely due to hydrophobic interactions. The MINDY3^EF-hand^ is not specific at binding to K48 chains as it can also bind to K11 and K63 homotypic and branched K48-K63 chains (**Fig. 3E**). To convincingly show that the MINDY3^EF-hand^ is a UBD, we mutated the proximal (site A) and distal (site C) Ub binding sites on the EF-hand. (**Fig. 3F**). Upon mutation of these sites, no binding between the mutant MINDY3^EF-hand^ and K48-Ub3 can be detected by either SEC or ITC (**Fig. 3I, EV3H**). In summary, these results show for the first time that an EF-hand domain can bind ubiquitin, establishing that the EF-hand domain of MINDY3 is a novel UBD that can bind to polyUb via simultaneous interactions with three Ub moieties.

### MINDY3 Interacts with RAD23A

The biological function of MINDY3 is poorly understood. Since our efforts to identify interactors of MINDY3 by immunoprecipitation-mass spectrometry approaches failed to identify significant binders, we performed a yeast two hybrid (Y2H) screen using FL MINDY3 as bait against a human placenta prey library. Thereby, we identified the proteasomal shuttling and nucleotide excision repair factor, RAD23A as a high-confidence interactor of MINDY3 (**Table 3**). RAD23A contains an N-terminal ubiquitin-like domain (UBL), RAD23A^UBL^, that interacts with the proteasome (Schauber *et al*, 1998), two ubiquitin associated (UBA) domains that preferentially bind K48-linked polyUb, and an XPC binding domain that binds to the XPC complex as part of the nucleotide excision repair process (**Fig. 4A**) (Raasi *et al*, 2004). To validate the Y2H screen result, we performed a pull-down experiment with recombinant GST-tagged RAD23A, which confirmed that purified MINDY3 directly binds to RAD23A (**Fig. 4B**). Furthermore, analysis by SEC reveals MINDY3 to form a stable complex with RAD23A **(Fig.4C**). Using ITC we quantified the binding affinity of the two proteins at 12.2 ± 4.22 μM (**Fig. 4D**). Of note, MINDY3 binds to Ub5 with 100-fold higher affinity compared to RAD23A. Furthermore, RAD23 binding results in an exothermic enthalpy change (ΔH = –4.84 ± 1.14 kJ/mol), whereas polyUb binding shows a strongly endothermic profile (ΔH = +11.0 ± 0.54 kJ/mol), indicative of an entropy-driven interaction dominated by hydrophobic effects and desolvation, suggesting that the binding mode and/or site(s) of RAD23A on MINDY3 are distinct from that of polyUb.

**Figure 4.**
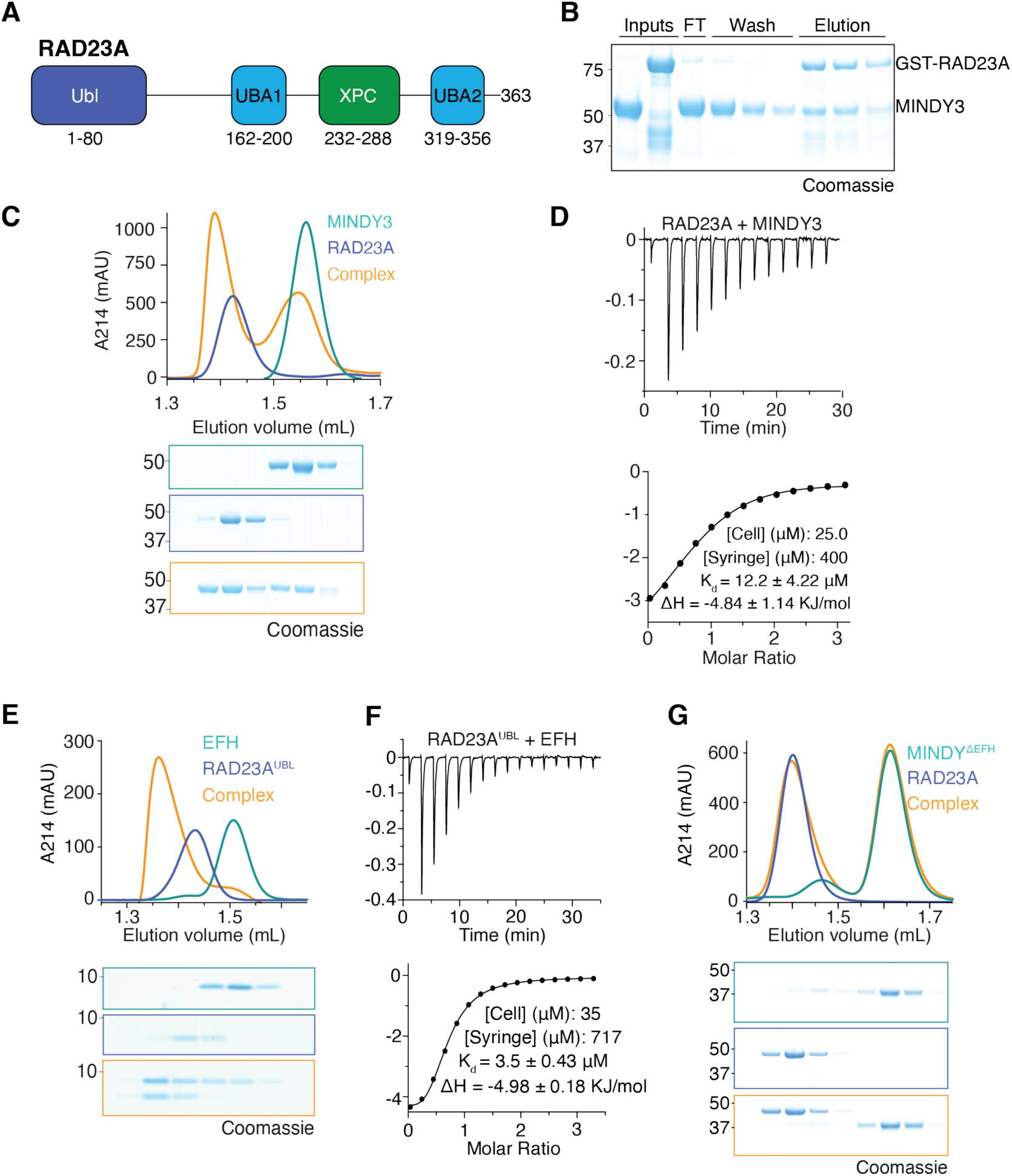
MINDY3 interacts with RAD23A. **(A)** Domain schematic of RAD23A showing the N-terminal UBL domain, the two UBA domains and the XPC-binding domain. **(B)** GST pull-down showing GST-RAD23A co-eluting with MINDY3. **(C)** Analytical SEC showing coelution of MINDY3 and RAD23A. **(D)** ITC experiments in which full-length RAD23A was titrated into full-length MINDY3. **(E)** Analytical SEC of MINDY3^EF-hand^ and RAD23A^UBL^ after pre-incubation. **(F)** ITC experiment in which the UBL of RAD23A was titrated into MINDY3^EF-hand^. **(G)** Analytical SEC of MINDY3^ΔEF-hand^ with RAD23A after pre-incubation.

To gain insight into how MINDY3 and RAD23A interact, we used AlphaFold to predict their binding mode (**Fig. EV4A-C**). In all predicted models of the FL MINDY3-RAD23A complex, the only region of RAD23A placed with high confidence is the UBL domain. This domain is modelled to interact with the MINDY3^EF-hand^ at site C, the putative proximal Ub binding site (**Fig. EV4B**). Indeed, the isolated MINDY3^EF-hand^ and the RAD23A^UBL^ domains coelute as a stable complex on SEC and the two domains bind with high affinity (Kd = 3.5 ± 0.43 μM) (**Fig. 4E, 4F**). As we could not detect binding of the MINDY3^EF-hand^ to monoUb, it is striking that it binds to the RAD23A^UBL^ domain with low micromolar affinity. Importantly, removal of the EF-hand in MINDY3 abolishes complex formation with RAD23A, confirming the MINDY3^EF-hand^ as the primary site of interaction with RAD23A (**Fig. 4G**).

Having discovered a direct interaction between MINDY3 and RAD23A by Y2H and subsequently confirmed this interaction *in vitro*, we next sought to check whether this interaction is also detectable in human cells. As RAD23B is very similar to RAD23A and has overlapping function, we also tested if MINDY3 can bind to RAD23B. Using transiently transfected FLAG-tagged FL or ΔEF-hand MINDY3 constructs, we were unable to co-immunoprecipitate (co-IP) RAD23A or RAD23B from either HEK293 or U2OS cells, confirming the unsuccessful CoIP-coupled mass spectrometry approach. Inferring that the interaction could be highly transient in cells, we repeated the experiment using the thiol-cleavable crosslinker DSP to stabilise temporarily formed complexes prior to FLAG IP. Under these conditions, we were able to detect an interaction of MINDY3 with both RAD23A and B in an EF-hand dependent manner (**Fig. EV4D**). We further corroborated this result by co-immunoprecipitating RAD23A using catalytically dead MINDY3^C51A^ in the absence of any crosslinking treatment (**Fig. EV4E**). Together, these results suggest that MINDY3 can interact with the RAD23 proteins in human cells via its EF-hand domain.

### Structure of MINDY3 EF-hand in complex with RAD23A UBL domain

To understand the interaction between RAD23A^UBL^ and MINDY3^EF-hand^ at the molecular level, we crystallised the reconstituted complex (**Fig. 5A**) and determined its high-resolution structure (**Table 2**). The RAD23A^UBL^ domain adopts the characteristic b-grasp fold observed in ubiquitin wherein five β-strands surround a central α-helix. The structure reveals that a hydrophobic ‘I49 patch’ on the UBL made up of Leu10, Ile49 and Val73 interacts with a hydrophobic surface on the EFH consisting of V338, Y343, L346 and the aliphatic portion of Lys350 (**Fig. 5B**). Of these residues, Val338, Tyr343 and Lys350 in MINDY3 are highly conserved throughout vertebrates. Individual Arginine substitutions of these interface residues, i.e. V338R, Y343R or L346R in MINDY3 disrupts complex formation with RAD23A, as shown by analytical SEC in which the two species now elute separately (**Fig. 5C**).

**Figure 5.**
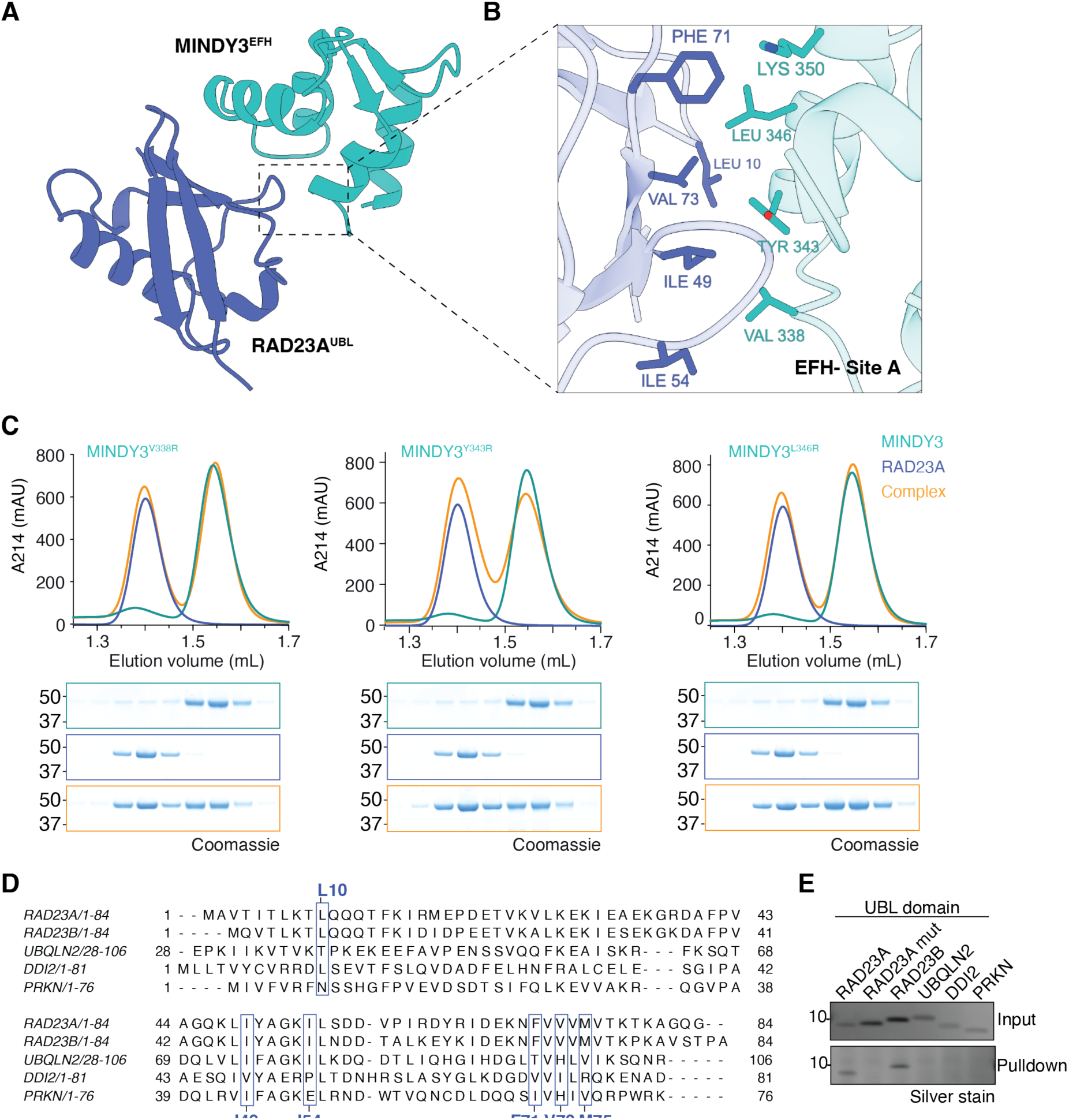
MINDY3^EFhand^ interacts specifically with RAD23A’s UBL domain. **A)** Crystal structure of MINDY3^EF-hand^ in complex with RAD23A^UBL^. **B)** Hydrophobic interactions between MINDY3^EF-hand^ and RAD23A^UBL^. **C)** SEC analyses using point mutants of MINDY3^EF-hand^ at the EFH-UBL interaction interface. **D)** Sequence alignment of UBL domains from several proteins. Residues involved in the interaction with MINDY3^EF-hand^ are highlighted. **E)** Pulldowns of HALO-MINDY3^EF-hand^ with a panel of UBLs.

**Table 2.**
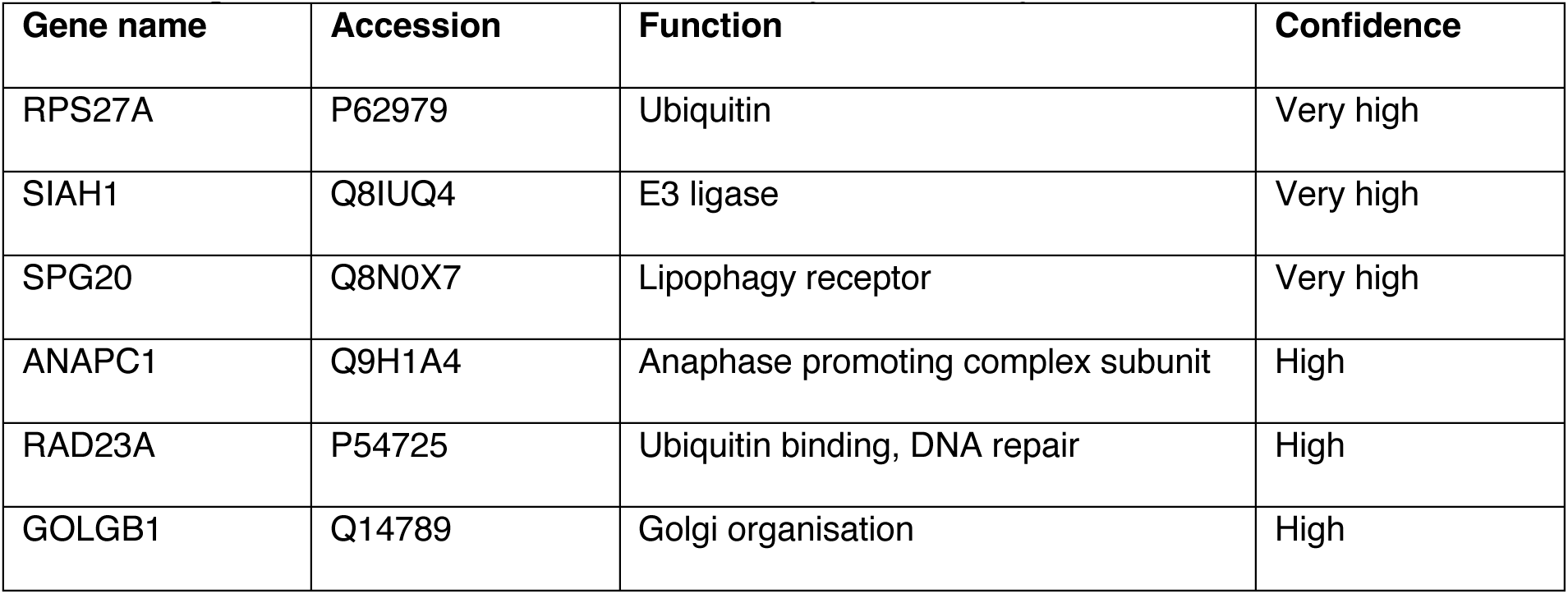
High confidence hits from MINDY3 yeast two-hybrid screen.

While the UBL of RAD23A only shares 32.1% identity and 56.0% similarity with ubiquitin, they adopt very similar structures. The ‘I49 patch’ on RAD23A^UBL^ shares similarities with the I44 patch of Ub, however, additional hydrophobic residues Met75, Phe71 and Ile54 of RAD23A^UBL^ mediate enhanced hydrophobic interactions (**Fig. EV5E**). Further hydrogen bonding is facilitated by Asn70, which interacts with Lys350 and Asp367. These additional interactions between the RAD23A^UBL^ and the MINDY3^EF-hand^ possibly explain why RAD23A^UBL^ binds to the MINDY3^EF-hand^ domain with high affinity, whilst no binding with monoUb was detected.

It was shown in different EF-hand domains that Ca^2+^ binding induces a conformational change from a ‘closed’ to an ‘open’ state. In the closed state, the helices adopt an almost antiparallel arrangement. Upon binding Ca^2+^, the top of the EF-hand is pinched together, which is accompanied by a shift in the helices to an open state in which a more perpendicular arrangement is adopted (Nelson & Chazin, 1998; Yap *et al*, 1999). This exposes formerly buried hydrophobic residues and creates a neo protein interaction site. In line with the lack of key metal-coordinating residues (**Fig. EV3B**), close inspection of the electron density map of the MINDY3^EF-hand^:RAD23A^UBL^ crystal structure did not reveal any additional density for Ca^2+^. The MINDY3^EF-hand^ adopts a conformation closer to the unbound, closed state (**Fig. EV5F**), and likely does not undergo any conformational transitions like canonical EF-hand domains.

To explore if the EF-hand specifically binds to the UBL domain of RAD23A, we tested binding of MINDY3^EF-hand^ to other UBLs. RAD23B^UBL^ shares 73.2% identity and 87.8% sequence similarity with RAD23A^UBL^, and importantly, the residues of the ‘I49 patch’ in RAD23A^UBL^ are conserved in RAD23B^UBL^ (**Fig. EV5B, C**). Many UBL domains analysed share similarities with RAD23A^UBL^ but lack several of the key interacting residues such as Leu10, Ile54, Phe71, Val73 and Met75 that make up the ‘I49 patch’. To determine whether the MINDY3^EF-hand^ is also able to bind other UBLs, we first assessed binding using SEC. While RAD23B^UBL^, like RAD23A^UBL^, forms a complex with the MINDY3^EF-hand^, PARKIN^UBL^ is unable to bind (**Fig. EV5D**), highlighting the importance of the ‘I49 patch’ in mediating UBL interactions within the EF-hand domain.

Next, we tested the binding of MINDY3^EF-hand^ to a panel of UBLs in an in vitro pulldown assay. While both RAD23A^UBL^ and RAD23B^UBL^ showed binding to the MINDY3^EF-hand^, we detected no interaction with UBLs from PARKIN (as above), DDI2 and UBQLN2, nor with RAD23A^UBL^ where key interacting residues Leu10, Phe71 and Val73 were mutated (**Fig. 5E**). Together these results suggest that MINDY3^EF-hand^ exclusively interacts with the RAD23A/B family UBL domain.

### MINDY3 is recruited to sites of DNA damage in a RAD23 dependent manner

In mammalian cells, RAD23 has two primary roles, namely facilitating DNA repair through the nucleotide excision repair (NER) pathway and shuttling ubiquitylated proteins to the proteasome for degradation (Grønbæk-Thygesen *et al*, 2023; Dantuma *et al*, 2009). To gain insights into the biological function of MINDY3 we assessed whether it could be implicated in either of these cellular pathways.

To determine whether MINDY3 plays a role in proteasomal shuttling, we generated stable cell lines expressing Ub^G76V^-GFP, a rapidly degraded ubiquitin fusion degron (UFD) protein (Dantuma *et al*, 2000). This reporter is degraded in a p97- and proteasome-dependent manner and has previously been reported as a substrate of RAD23 (Gödderz *et al*, 2017). We assessed its stability by immunofluorescence and western blotting, however, we found no major effect on the degradation of Ub^G76V^-GFP (**Fig. EV6A-D**), suggesting that MINDY3 does not strongly contribute to RAD23-mediated proteasomal shuttling.

To explore whether MINDY3 is implicated in DNA damage repair, we first assessed whether MINDY3 is recruited to sites of DNA damage. To this end, we generated U2OS cells stably expressing GFP-tagged MINDY3 and monitored its localisation following micro-irradiation (microIR). Intriguingly, MINDY3 was found to be rapidly recruited to sites of DNA damage (**Fig. 6A**). Strikingly, we observed that this recruitment of MINDY3 requires the EF-hand domain, as FL GFP-MINDY3 was recruited to DNA damage sites in 34% of irradiated cells, but absence of the EF-hand domain fully abrogated this recruitment (**Fig. 6B, C**).

**Figure 6.**
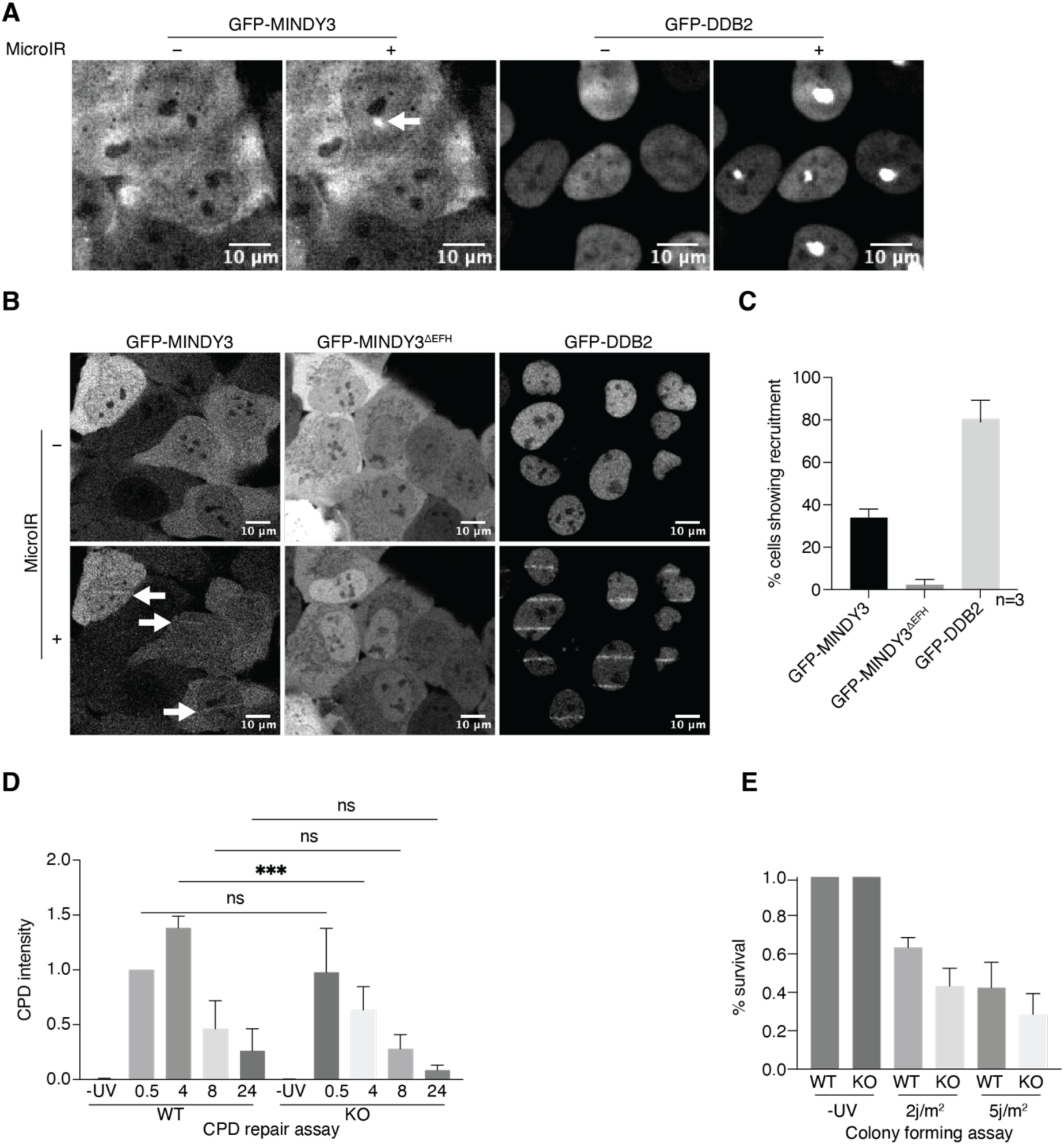
MINDY3 is recruited to sites of DNA damage in a RAD23A dependent manner. **A)** Representative images of microIR U2OS cells stably expressing GFP-MINDY3 and GFP-DDB2. **B)** Representative images of U2OS cells expressing GFP-DDB2, GFP-MINDY3 and GFP-MINDY3^DEF-hand^ after microIR. **C)** Quantification of the percentage of cells in which recruitment of the specified protein to DNA damage sites was observed. Quantification was performed on three independent experiments with 20-50 cells counted for each experiment. **D)** U2OS, WT and MINDY3 KO cells were irradiated with UV-C (20 J/m^2^), CPD abundance was monitored by dot blotting genomic DNA extracted at the indicated times and probing with a CPD and CDNA antibody. Quantification was performed on three biological replicates. CPD signal was adjusted based on the cDNA signal and values were normalised to WT time 0.5 h. **E)** Quantification of colonies formed by U2OS cells, WT and MINDY3 KO after UV-C irradiation with the indicated doses. Quantification of four independent experiments performed in technical triplicates is shown.

We next investigated if the recruitment of MINDY3 to DNA damage sites correlated with a role in DNA repair. As RAD23 plays a role in NER, the primary repair pathway for UV-damaged DNA, we assessed if NER is defective in the absence of MINDY3. To this end we assessed the clearance of UV induced cyclobutane pyrimidine dimers (CPDs) by DNA dot blot assays. A difference in the abundance of CPD lesions 4h after irradiation was observed in MINDY3 KO cells, importantly however similar levels of CPD lesions were observed in WT and MINDY3 KO cells at 8 and 24 h indicating that repair of CPD lesions was not defective in the absence of MINDY3 (**Fig. 6D**). Furthermore, we observed using colony forming assays that MINDY3 deficiency did not robustly affect cell survival after UV irradiation (**Fig. 6E**). These experiments suggest that the absence of MINDY3 does not lead to a defective NER pathway.

### MINDY3 cleaves ubiquitin from RAD23-bound substrates

Having shown a *bona fide* direct interaction between MINDY3^EF-hand^ and RAD23A^UBL^ in both in vitro and in human cells, we next assessed the effect of this interaction on the DUB activity of MINDY3. One possibility is that RAD23A binding to the MINDY3^EF-hand^ domain affects MINDY3 DUB activity. Therefore, MINDY3 was pre-incubated with RAD23A in 1:1 and 1:2 ratios to pre-form the complex. K48-Ub_5_ was added, and its cleavage was monitored in the established DUB assay (**Fig. 7A**). After 60 minutes, similar amounts of lower MW cleavage products are formed even when MINDY3 is complexed with RAD23A. Hence, the binding of RAD23A to MINDY3 does not compromise or promote its ability to cleave polyUb chains.

**Figure 7.**
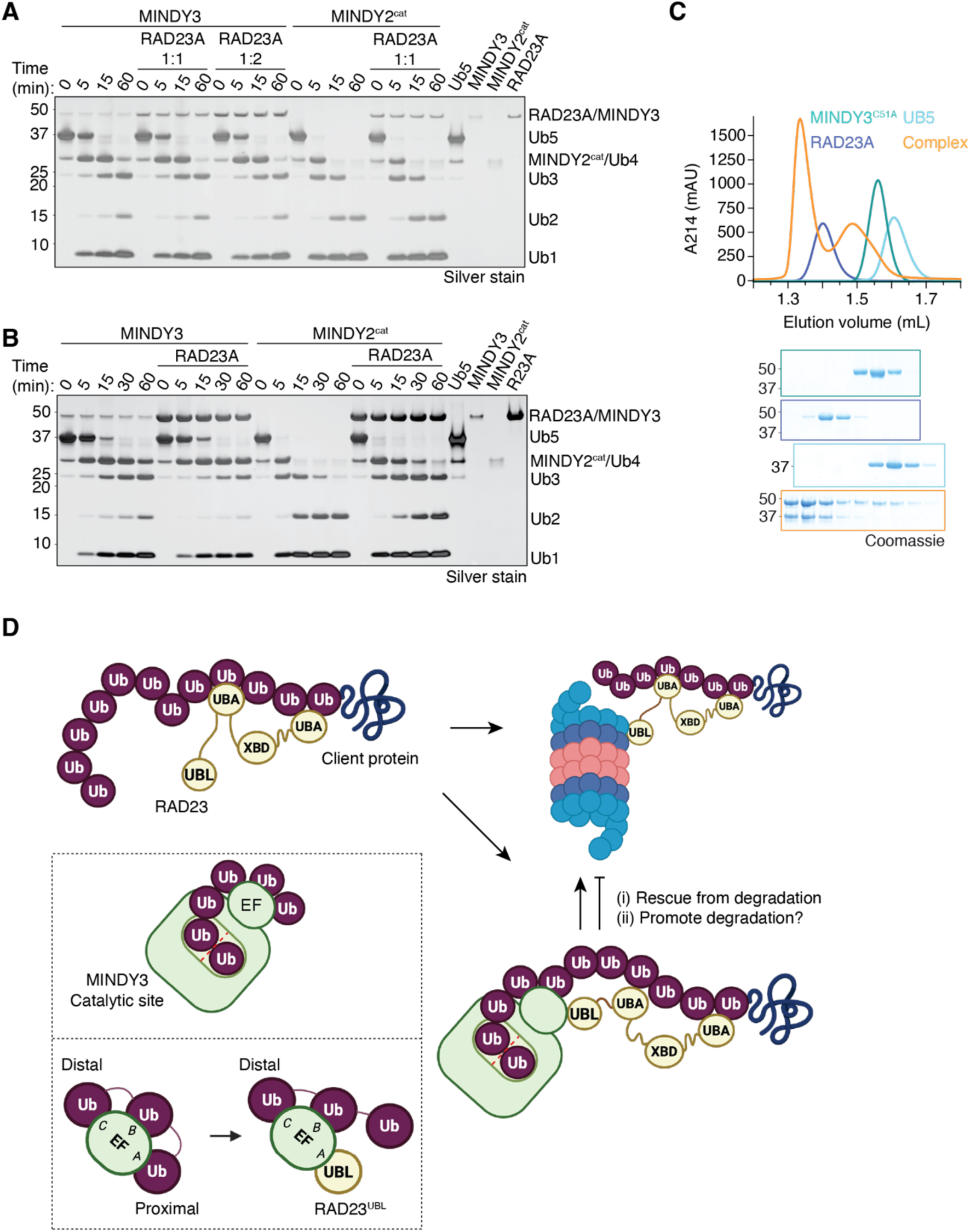
MINDY3 can cleave ubiquitin chains bound by RAD23A. **A)** DUB assay in which MINDY3 FL or MINDY2 catalytic domain (cat) was incubated with increasing concentrations of RAD23A prior to the addition of K48-linked Ub5. **B)** DUB assay with MINDY3 in which K48-Ub5 was pre-incubated with RAD23A. **C)** Analytical SEC of MINDY3^C51A^, K48-Ub5 and RAD23A. **D)** Schematic model showing the role of RAD23A in proteasomal shuttling positing the role of MINDY3 in this process.

We next investigated whether RAD23A which can bind to K48-linked chains through its UBA domains protects ubiquitin chains from cleavage by hindering DUB access. To this end, we first complexed RAD23A with polyUb chains to mimic a situation where RAD23A is bound to ubiquitylated client and investigated if MINDY3 can still cleave these RAD23A-bound chains. (**Fig. 7B**). After 60 minutes, cleavage of polyUb within pre-formed RAD23A-Ub5 by MINDY3 only has a minor effect on MINDY3 cleavage of chains. In contrast, when the same experiment was performed using MINDY2, which does not bind RAD23A, a pre-formed RAD23A-Ub5 complex has a drastic inhibitory effect on cleavage by MINDY2 compared to Ub5 alone (**Fig. 7B**). MINDY3 is therefore able to efficiently cleave polyUb chains even when they are bound by RAD23A. This suggests that formation of a ternary complex of MINDY3:RAD23A:K48-Ub5 is possible. Indeed, SEC using catalytically dead MINDY3 confirms the formation of such a trimeric complex (**Fig. 7C**). Further, AlphaFold modelling of a MINDY3:RAD23A:K48-Ub5 complex shows that the RAD23^UBL^ domain binds to Site C on the MINDY3^EF-hand^ domain that forms the S4’ site, thus displacing the S4 ubiquitin and therefore would not affect DUB activity (see below). Hence, MINDY3 could edit Ub chains present on RAD23 clients.

## Discussion and future perspectives

Our work reveals that MINDY3 like MINDY1/2 prefers to cleave long ubiquitin chains. However, we find that MINDY3 uses a distinct mechanism as it contains a unique ubiquitin binding EF-hand domain. Alphafold modelling of MINDY3 with multiple ubiquitin molecules produces a model in which four ubiquitin molecules are placed in positions and in orientations that could represent how MINDY3 binds polyUb **(Fig. EV7A, B)**. While the distal ubiquitin is placed at the S1 site, the proximal ubiquitin is not placed at the S1’ site. Some structural rearrangement may be required to form the S1’ site. The third ubiquitin in this hypothetical chain is then placed at the S2’ site. Based on the geometry of ubiquitin, the positioning of the K48 of the S2’ Ub and the G76 of the S1 Ub, the approximate location of the S1’ site can be inferred. The MINDY3^EF-hand^ acts as a UBD and can bind 3 ubiquitin moieties. AlphaFold places ubiquitin moieties on the EF-hand at the S3’ (site A) and at the UBL binding S5’ site (site C) (using notation from **Fig. 3D**). Overlaying the model of the EF-hand as a UBD (**Fig. 3D**) with the model in **Fig EV7A** results in severe clashing between the medial ubiquitin at site B (the S4’ ubiquitin) and the S1 ubiquitin, which explains why AlphaFold was unable to place a ubiquitin at this site. However, the MINDY3^EF-hand^ is attached to the core catalytic domain of MINDY3 by two flexible linkers and so the positioning of the MINDY3^EF-hand^ could vary. Some movement in the MINDY3^EF-hand^ could therefore allow the medial ubiquitin to be accommodated at the S4’ site (site B MINDY3^EF-hand^). AlphaFold predictions using MINDY3 can therefore place four putative ubiquitin binding sites on MINDY3 but can be used to infer the existence of six ubiquitin binding sites **(Fig. EV7C)**. It is remarkable that despite the divergence in sequence and structural similarity between MINDY3 and MINDY1/2, they show selectivity not only for cleaving K48-linkages but also long polyUb chains, albeit through distinct mechanisms.

RAD23A/B have also been shown previously to interact with DUBs. For instance, ATXN3 has two known ubiquitin binding sites on its catalytic domain, namely S1 and S2 (Weeks *et al*, 2011; Nicastro *et al*, 2009, 2010). ATXN3, like MINDY3, prefers cleaving longer and branched polyUb chains, but prefers cleaving K63-linkages (Winborn *et al*, 2008; Lange *et al*, 2024). In addition to binding ubiquitin, the S2 site can also bind RAD23A/B via their UBL domains (Wang *et al*, 2000; Nicastro *et al*, 2010, 2009). Disruption of this interaction by mutating the S2 site results in a marked increase in its degradation by the proteasome, whilst mutation of the polyUb binding UIMs of ATXN3 opposes this degradation (Blount *et al*, 2014).

The precise functional relationship between MINDY3 and RAD23A/B remains to be uncovered. There are several scenarios that can be envisioned: i) RAD23A/B shuttles MINDY3 to the proteasome for degradation ii) RAD23A/B binding stabilises MINDY3 levels as with ATXN3, iii) MINDY3 binds to and deubiquitylates polyUb cargo bound to RAD23A/B thereby rescuing the cargo from proteasomal degradation, iv) MINDY3 is shuttled to sites of DNA damage by RAD23A/B where it then deubiquitylates targets (**Fig. 7D**). Preliminary data shows that knockout or knockdown of MINDY3 does not affect abundance of RAD23s or, vice versa (**Fig. EV7D**), suggesting that the latter two options may be more likely.

Our work suggests that RAD23A/B bound to ubiquitin chains can dock onto MINDY3 via the UBL domain:EF-hand interaction, and lead to the editing or removal of ubiquitin chains on the RAD23A/B client. The UBL domains of human RAD23A and B primarily interact with the proteasome via UIM2 of RPN10, with affinities in the low µM range (Mueller & Feigon, 2003; Fujiwara *et al*, 2004). This binding affinity is comparable to that of the RAD23A^UBL^ for MINDY3^EF-hand^. Notably, regions of RAD23A/B outside the UBL have no effect on binding to RPN10 or to MINDY3 (Hiyama *et al*, 1999) (**Fig. 4**). This similarity in binding affinities of RAD23A with either RPN10 or MINDY3 suggests that some competition between the two proteins for RAD23A (or B) binding could be envisioned (**Fig. 7D**).

Future work focussing on exploring these possibilities and identifying the targets of MINDY3 deubiquitylation will shed light on the physiological importance of this interaction. However, a major challenge to establishing the role of MINDY3 in regulating the ubiquitylation status and fate of RAD23-bound clients is that the identities of the physiological proteasomal clients of RAD23A/B are unknown. Similarly, identifying the targets of MINDY3 at DNA damage sites will be of great interest. Here, redundancy or overlapping functions with other DUBs may make it daunting to identify the precise substrates of MINDY3 and how it regulates DNA damage responses.

## Materials and Methods

### Methods and Protocols

#### Reagents used in this study

A list of plasmids used in this study can be found in **Table 3**. All DNA constructs used were generated by L.A.A., M.R.M., R.K., and the cloning team at the Medical Research Council Protein Phosphorylation and Ubiquitylation Unit (MRC PPU) Reagents and Services. All plasmids are available upon request by contacting Y.K. or at https://mrcppureagents.dundee.ac.uk/. A list of siRNA sequences are listed in **Table 4**.

**Table 3.**
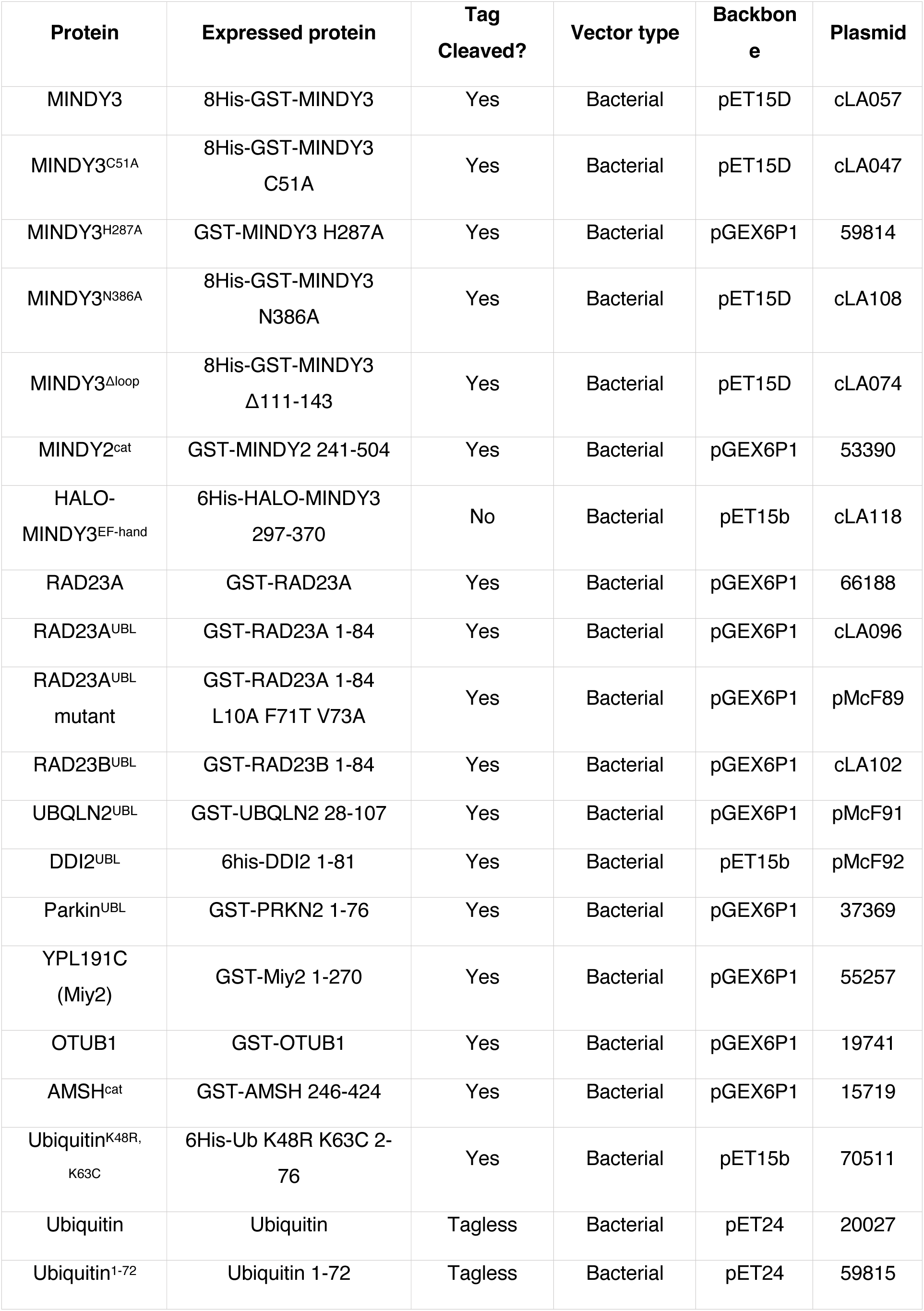
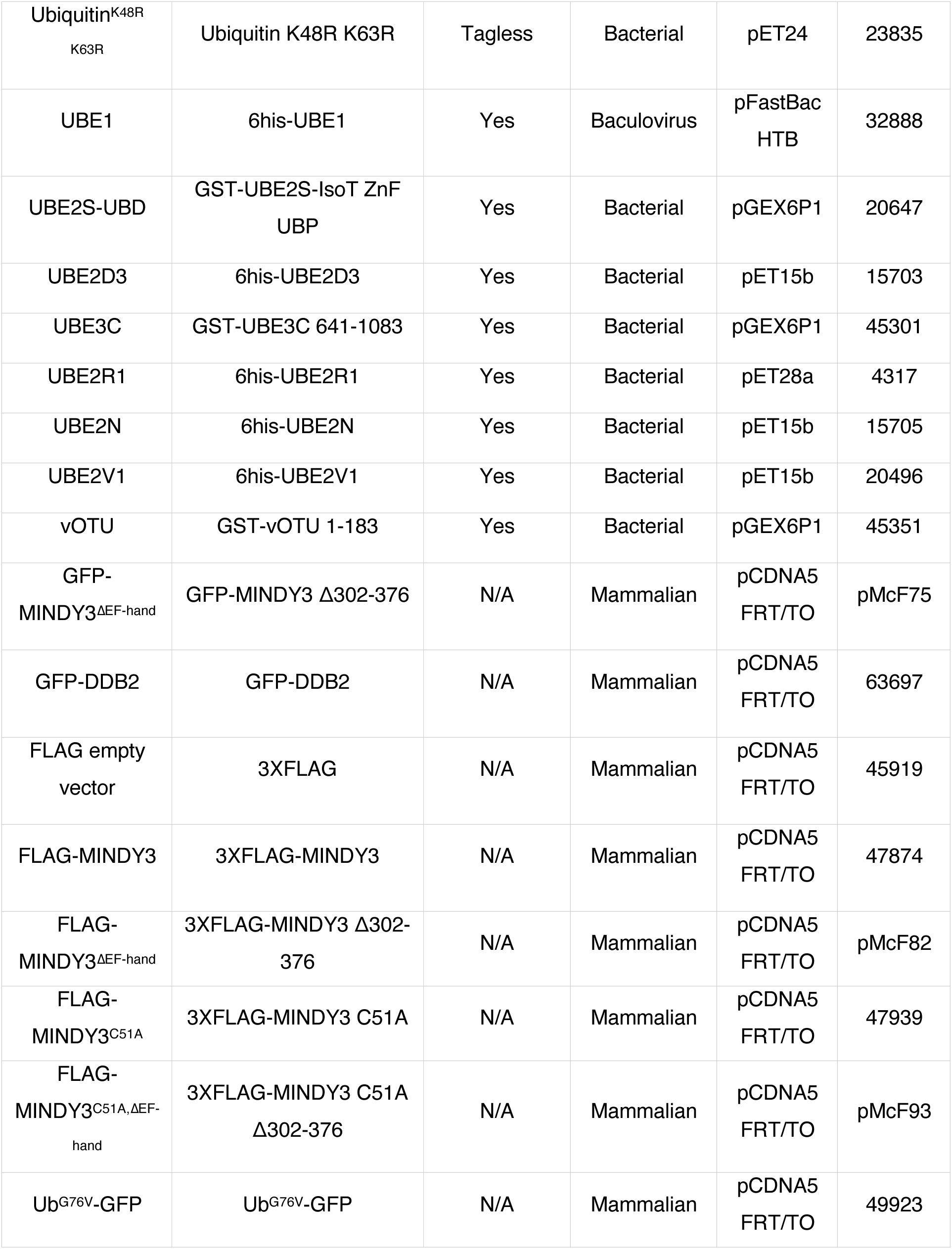
Plasmids used in this study.

| Protein | Expressed protein | Tag Cleaved? | Vector type | Backbone | Plasmid |
| --- | --- | --- | --- | --- | --- |
| MINDY3 | 8His-GST-MINDY3 | Yes | Bacterial | pET15D | cLA057 |
| MINDY3<sup>C51A</sup> | 8His-GST-MINDY3<br>C51A | Yes | Bacterial | pET15D | cLA047 |
| MINDY3<sup>H287A</sup> | GST-MINDY3 H287A | Yes | Bacterial | pGEX6P1 | 59814 |
| MINDY3<sup>N386A</sup> | 8His-GST-MINDY3<br>N386A | Yes | Bacterial | pET15D | cLA108 |
| MINDY3<sup>Δloop</sup> | 8His-GST-MINDY3<br>Δ111-143 | Yes | Bacterial | pET15D | cLA074 |
| MINDY2<sup>cat</sup> | GST-MINDY2 241-504 | Yes | Bacterial | pGEX6P1 | 53390 |
| HALO-MINDY3<sup>EF-hand</sup> | 6His-HALO-MINDY3<br>297-370 | No | Bacterial | pET15b | cLA118 |
| RAD23A | GST-RAD23A | Yes | Bacterial | pGEX6P1 | 66188 |
| RAD23A<sup>UBL</sup> | GST-RAD23A 1-84 | Yes | Bacterial | pGEX6P1 | cLA096 |
| RAD23A<sup>UBL</sup><br>mutant | GST-RAD23A 1-84<br>L10A F71T V73A | Yes | Bacterial | pGEX6P1 | pMcF89 |
| RAD23B<sup>UBL</sup> | GST-RAD23B 1-84 | Yes | Bacterial | pGEX6P1 | cLA102 |
| UBQLN2<sup>UBL</sup> | GST-UBQLN2 28-107 | Yes | Bacterial | pGEX6P1 | pMcF91 |
| DDI2<sup>UBL</sup> | 6his-DDI2 1-81 | Yes | Bacterial | pET15b | pMcF92 |
| Parkin<sup>UBL</sup> | GST-PRKN2 1-76 | Yes | Bacterial | pGEX6P1 | 37369 |
| YPL191C<br>(Miy2) | GST-Miy2 1-270 | Yes | Bacterial | pGEX6P1 | 55257 |
| OTUB1 | GST-OTUB1 | Yes | Bacterial | pGEX6P1 | 19741 |
| AMSH<sup>cat</sup> | GST-AMSH 246-424 | Yes | Bacterial | pGEX6P1 | 15719 |
| Ubiquitin<sup>K48R,</sup><br><sup>K63C</sup> | 6His-Ub K48R K63C 2-76 | Yes | Bacterial | pET15b | 70511 |
| Ubiquitin | Ubiquitin | Tagless | Bacterial | pET24 | 20027 |
| Ubiquitin<sup>1-72</sup> | Ubiquitin 1-72 | Tagless | Bacterial | pET24 | 59815 |
| Ubiquitin<sup>K48R</sup><br>K63R | Ubiquitin K48R K63R | Tagless | Bacterial | pET24 | 23835 |
| UBE1 | 6his-UBE1 | Yes | Baculovirus | pFastBac<br>HTB | 32888 |
| UBE2S-UBD | GST-UBE2S-IsoT ZnF<br>UBP | Yes | Bacterial | pGEX6P1 | 20647 |
| UBE2D3 | 6his-UBE2D3 | Yes | Bacterial | pET15b | 15703 |
| UBE3C | GST-UBE3C 641-1083 | Yes | Bacterial | pGEX6P1 | 45301 |
| UBE2R1 | 6his-UBE2R1 | Yes | Bacterial | pET28a | 4317 |
| UBE2N | 6his-UBE2N | Yes | Bacterial | pET15b | 15705 |
| UBE2V1 | 6his-UBE2V1 | Yes | Bacterial | pET15b | 20496 |
| vOTU | GST-vOTU 1-183 | Yes | Bacterial | pGEX6P1 | 45351 |
| GFP-<br>MINDY3<sup>ΔEF-hand</sup> | GFP-MINDY3 Δ302-376 | N/A | Mammalian | pCDNA5<br>FRT/TO | pMcF75 |
| GFP-DDB2 | GFP-DDB2 | N/A | Mammalian | pCDNA5<br>FRT/TO | 63697 |
| FLAG empty<br>vector | 3XFLAG | N/A | Mammalian | pCDNA5<br>FRT/TO | 45919 |
| FLAG-MINDY3 | 3XFLAG-MINDY3 | N/A | Mammalian | pCDNA5<br>FRT/TO | 47874 |
| FLAG-<br>MINDY3<sup>ΔEF-hand</sup> | 3XFLAG-MINDY3 Δ302-376 | N/A | Mammalian | pCDNA5<br>FRT/TO | pMcF82 |
| FLAG-<br>MINDY3<sup>C51A</sup> | 3XFLAG-MINDY3 C51A | N/A | Mammalian | pCDNA5<br>FRT/TO | 47939 |
| FLAG-<br>MINDY3<sup>C51A,ΔEF-</sup><br><sup>hand</sup> | 3XFLAG-MINDY3 C51A<br>Δ302-376 | N/A | Mammalian | pCDNA5<br>FRT/TO | pMcF93 |
| Ub<sup>G76V</sup>-GFP | Ub<sup>G76V</sup>-GFP | N/A | Mammalian | pCDNA5<br>FRT/TO | 49923 |

**Table 4.**
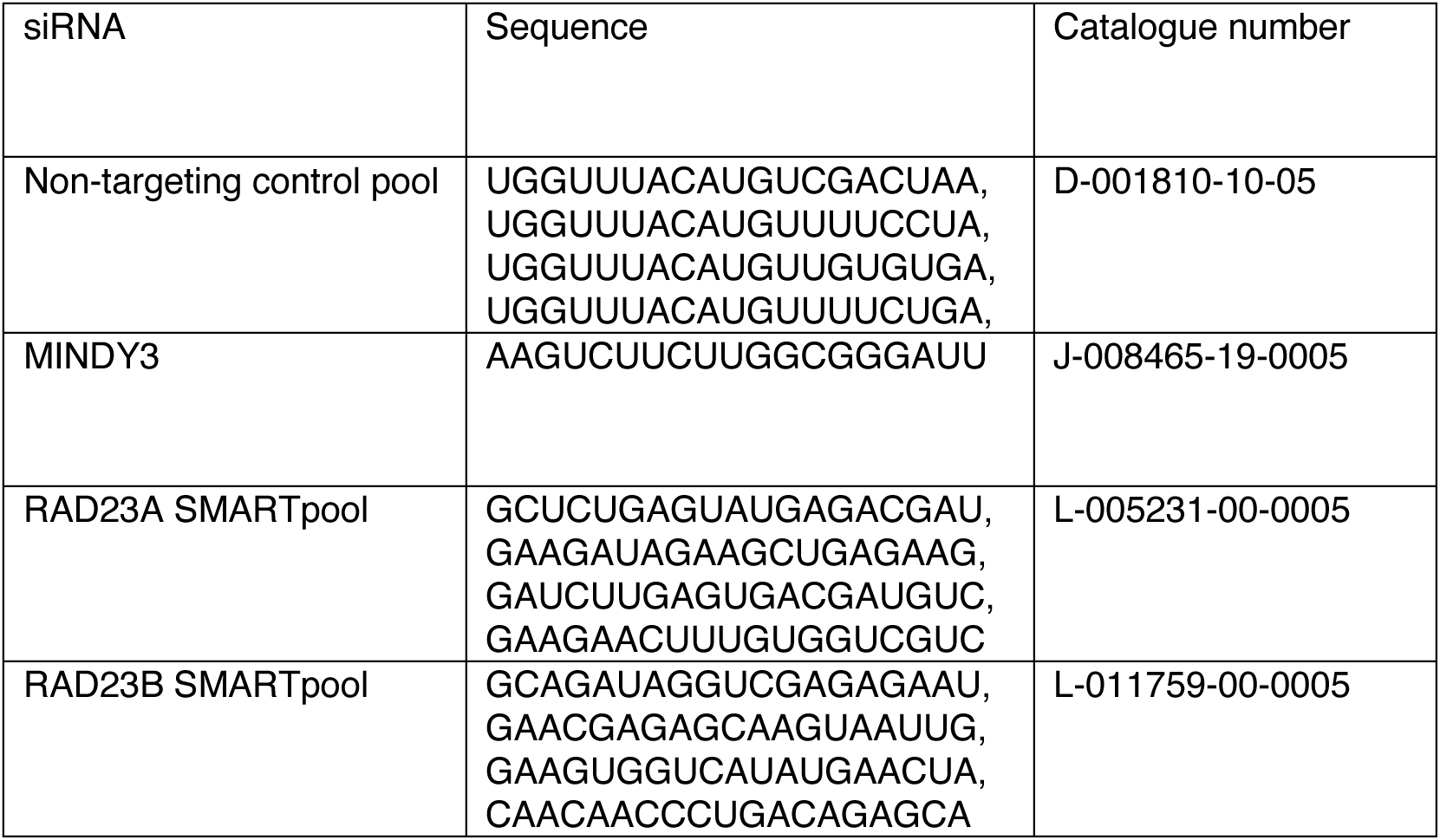
siRNAs used in this study.

##### Protein Expression

cDNA was transformed into E. coli BL21 cells and the transformant mixture was streaked onto LB agar plates supplemented with either 100 μg/mL ampicillin or 50 μg/mL kanamycin. After incubation overnight at 37°C, a single colony was picked and used to inoculate 100 mL 2x TY medium. Alternatively, a glycerol stock was used for inoculation. The culture was grown at 37°C/180rpm until an OD600 of 0.6-0.8 was reached. The temperature of the incubator was then lowered to 18°C and left to acclimatise for 30 mins. Protein expression was then induced by the addition of IPTG to a final concentration of 300 μM and further incubation at 18°C for 18 hrs. For the expression of untagged monoUb (and derivative point mutants), expression was induced with 1 mM IPTG for 5.5 hrs at 37°C. Cultures were then harvested by centrifugation at 4200 rpm for 30 mins at 4°C. The supernatant was discarded, and the pellet resuspended in an appropriate lysis buffer. The pellet was then flash-frozen in liquid nitrogen and stored at -80°C until required.

##### Protein purification

Pellets of cells were thawed in a water bath at 37°C and then incubated on ice for 15 mins. E. coli cultures were lysed by sonication. Cell lysate was then clarified by centrifugation at 35,000g for 30 mins at 4°C. The clarified lysate was then filtered through a fine mesh (BRAND) to remove any insoluble material.

###### GST-tagged

For GST-tagged proteins, the lysate was incubated with pre-equilibrated Glutathione Sepharose 4B resin for 1 hour at 4°C. The resin was then washed with 100 bed volumes of GST high salt buffer followed by 50 bed volumes of GST low salt buffer. For tag removal, the resin was then incubated o/n at 4°C with C3 protease to cleave the C3 site between the protein and the GST tag. Alternatively, GST-tagged recombinant protein was eluted from the resin by the addition of LSB supplemented with 30 mM reduced glutathione. Eluted protein was concentrated using an Amicon Ultra Centrifugal Filter, the purity assessed by SDS-PAGE, flash-frozen and stored at -80°C.

###### His-tagged

For 6-10His-tagged proteins, the lysate was incubated with pre-equilibrated His60 Superflow resin (TaKaRa) and incubated at 4°C for 1 hour. The resin was then washed with 20 bed volumes binding buffer, 20 bed volumes wash buffer, and finally eluted in 10 bed volumes of elution buffer. Eluted protein was buffer exchanged to storage buffer, concentrated using an Amicon Ultra Centrifugal Filter, the purity assessed by SDS-PAGE, flash-frozen and stored at -80°C.

###### 8His-GST-tagged

For proteins that were His-GST tagged, the protocol for His-tag purification described above was followed. Protein was eluted from the His60 Superflow resin and the eluent was then incubated with pre-equilibrated Glutathione Sepharose as described above.

###### Untagged monoUb

50 mM sodium acetate pH 4.5 was added to the clarified lysate to precipitate host proteins. This was incubated at RT overnight on a roller. The precipitate was then pelleted by centrifugation at 4200 rpm for 30 mins at 4°C. The clarified supernatant was decanted, filtered through a 0.45 μM membrane filter and ubiquitin was purified by cation exchange chromatography. Fractions were analysed by SDS-PAGE, pooled, concentrated using Amicon Ultra Centrifugal Filters and buffer exchanged into 50 mM Tris-HCl pH 7.5. Protein was then flash-frozen and stored at -80°C.

###### Ion exchange chromatography

After affinity chromatography, some proteins were further purified by ion exchange chromatography. For anion exchange chromatography, protein was passed over a ResourceQ 6 mL column (Cytiva). The column was washed with Buffer A (30 mM NaCl, 50 mM Tris-HCl pH 8.5) until the UV_280_ baseline stabilised. Protein was then eluted in a gradient of Buffer B (1000 mM NaCl, 50 mM Tris-HCl pH 8.5). For ubiquitin and polyUb purification by cation exchange chromatography, protein was passed over a ResourceS 6 mL column (Cytiva). The column was washed with Buffer A (50 mM sodium acetate pH 4.5) until the UV_214_ baseline stabilised (ubiquitin only weakly absorbs at UV_280_). Protein was then eluted in a gradient of Buffer B (1000 mM NaCl, 50 mM sodium acetate pH 4.5) with increasing length of polyUb chain eluting at an increased % of buffer B. Fractions were analysed by SDS-PAGE.

###### Size exclusion chromatography

Protein was concentrated to an appropriate volume for the selected loop – usually a 1 mL loop. Protein was then filtered through a Costar Spin-X centrifuge filter (Corning) and injected into the loop. Protein was then loaded onto either a Superdex 75 16/600 or Superdex 200 16/600 column pre-equilibrated with GFC buffer. Fractions were analysed by SDS-PAGE, pooled and concentrated using Amicon Ultra Centrifugal Filters. Protein was then flash-frozen and stored at -80°C.

###### MINDY3apo expression and purification for crystallisation

The ORF of human MINDY3 was cloned into the pET-M30 bacterial expression vector using NcoI and XhoI restriction sites. The vector encoded an N-terminal 6×His tag, GST tag, and a TEV protease cleavage site. Expression was performed in E. coli pRARE cells grown in LB medium supplemented with chloramphenicol and kanamycin until reaching an OD₆₀₀ of ∼0.7. For selenomethionine (SeMet) labeling, cultures were grown in M9 minimal medium using a SelenoMet Medium Kit (Molecular Dimensions) according to the manufacturer’s instructions. Upon reaching OD₆₀₀ ∼0.7, cultures were cooled to 18 °C, induced with 1 mM IPTG, and incubated overnight for protein expression. Cells were harvested by centrifugation at 9000 × g for 10 min, and cell pellets were stored at –80 °C until further use. Pellets were resuspended in ice-cold lysis buffer (50 mM Tris pH 7.5, 300 mM NaCl, 5% (v/v) glycerol, 10 mM imidazole, 2 mM MgCl₂, 1 mM β-mercaptoethanol) and lysed by sonication. The lysate was clarified by centrifugation at 60,000 × g for 45 min, and the supernatant was applied to Ni-NTA agarose resin pre-equilibrated with lysis buffer. Following an initial 10 CV wash with lysis buffer, the resin was washed with 5 CV of lysis buffer containing 1 M NaCl, and the protein was eluted with lysis buffer containing 250 mM imidazole. The eluate was loaded onto a GSTprep column (Cytiva) pre-equilibrated with lysis buffer. After 2 h of incubation, the column was washed, and bound protein was eluted using buffer containing 50 mM Tris pH 7.5, 300 mM NaCl, 2 mM DTT, and 18 mM reduced glutathione. Affinity tags were cleaved off by 6×His-tagged TEV during an overnight dialysis at 4 °C against the dialysis buffer (20 mM Tris pH 7.5, 150 mM NaCl, 2 mM DTT). Cleaved tags and TEV protease were removed by subsequent reverse-affinity Ni-NTA and GST columns. The flowthrough was concentrated and subjected to size-exclusion chromatography (SEC) using a HiLoad 16/600 Superdex 75 column (Cytiva) equilibrated in 20 mM Tris pH 7.5, 150 mM NaCl, and 2 mM DTT. Fractions corresponding to monomeric MINDY-3 were pooled, concentrated, and either used immediately or snap-frozen in liquid nitrogen and stored at –80 °C.

##### Polyubiquitin chain assembly & purification

###### K11-linked polyUb

K11-linked chains were assembled in chain ligation buffer containing 1500 μM Ub, 1 μM UBE1, 40 μM UBE2S-UBD. This reaction also produces a small quantity of K63-linked chains, so the mix is supplemented with 2 μM AMSH to cleave them. The reaction was incubated at 30°C for 6 hrs. 2 μM of fresh AMSH was then added along with 5 μM DTT and the reaction was incubated for a further 16 h.

###### K29-linked polyUb

K29-linked chains were assembled in chain ligation buffer containing 1500 μM Ub, 0.64 UBE1, 9.5 μM UBE2D3 and 3 μM UBE3C. The reaction was incubated at 30°C for 6 h. This reaction also produces a variety of linkage types, so the reaction is then supplemented with vOTU and 5 μM DTT, and incubated overnight at 30°C.

###### K48-linked polyUb

K48-linked chains were assembled in chain ligation buffer containing 1500 μM Ub, 1 μM UBE1, 25 μM UBE2R1. The reaction was incubated for 6 hours at 30°C. For unlabelled longer fluorescent chains, the reaction was incubated for 18 h at 30°C.

###### K63-linked polyUb

K63-linked chains were assembled in chain ligation buffer containing 1500 μM Ub, 1 μM UBE1, 10 μM UBE2N and 20 μM UBE2V1. The reaction was incubated for 6 hours at 30°C.

###### Branched K48-K63 linked polyUb

Branched K48–K63 trimer was assembled in two steps. First, K63-linked dimer was assembled in chain ligation buffer containing 1000 µM Ub^1-72^, 1500 µM Ub^K48R,K63R^, 1 µM UBE1, 20 µM UBE2N and 20 µM UBE2V1. Reaction was incubated at 30°C for 6 hours. Second, K48 branch was added in chain ligation buffer containing 350 µM step 1 product, 700 µM Ub^K48R,K63R^, 1 µM UBE1, and 25 µM UBE2R1. Reaction was incubated at 30°C for 6 hours.

###### Purification of homotypic chains

The reaction mixes were quenched by the addition of 50 mM sodium acetate pH 4.5. The reaction mix was then filtered through a 0.45 micron filter (Sartorius). The heterogeneous mix of polyUb chains were then separated by cation exchange chromatography. The mix was passed over a ResourceS 6 mL column (Cytiva). The column was washed with Buffer A (50 mM sodium acetate pH 4.5) until the UV_214_ baseline stabilised (ubiquitin only weakly absorbs at UV_280_). Protein was then eluted in a gradient of Buffer B (1000 mM NaCl, 50 mM sodium acetate pH 4.5) with increasing length of polyUb chain eluting at an increased % of buffer B. Fractions were analysed by SDS-PAGE.

###### Fluorescent labelling of K48-linked polyUb chains

Fluorescently labelled Ub2-Ub5 were synthesised as described previously (Rehman *et al*., 2016).

###### Assembly of K48-Ub2 containing Cys residue at the Ub^dist^

*(K48-Ub2 [Cys-Ub_dist_])* K48-Ub2 [Cys-Ub_dis_] was assembled enzymatically using Ub G75A/G76A as the Ub acceptor and Ub K48R with Cys residue upstream of M1 (Cys-Ub K48R) as the Ub donor. The assembly reaction was carried out for 1 h at 30 °C in 1 ml reaction containing 1 mM Ub G75A/G76A, 1 mM Cys-Ub K48R, 0.5 μM UBE1, 15 μM UBE2R1, 10 mM ATP, 50 mM Tris-HCl (pH 7.5), 10 mM MgCl2, and 0.6 mM DTT.

###### Assembly of K48-Ub3 containing Cys residue at the Ub^dist^ (K48-Ub3 [Cys-Ub_dist_])

K48-Ub3 [Cys-Ub_dist_] was assembled by capping a pre-assembled K48-Ub2 by Cys-Ub K48R (donor Ub). The assembly reaction was carried out for 2 h at 30 °C in 1 ml reaction containing 250 μM K48-Ub2, 1.25 mM Cys-Ub K48R, 0.5 μM UBE1, 15 μM UBE2R1, 10 mM ATP, 50 mM Tris-HCl (pH 7.5), 10 mM MgCl2, and 0.6 mM DTT. K48-Ub3 [Cys-Ub_dis_] was purified as described previously (Kristariyanto *et al*, 2015).

###### Assembly of K48-Ub5 containing Cys residue at the Ub^prox^ (K48-Ub5 [Cys-Ub_prox_])

K48-Ub5 [Cys-Ub_prox_] was assembled by extending Ub 1-75 (Ub acceptor), which contains Cys residue upstream of M1 (Cys-Ub 1-75), with a pre-assembled K48-Ub2. The assembly reaction was carried out for 1.5 h at 30 °C in 1 ml reaction containing 250 μM K48-Ub2, 1.25 mM Cys-Ub 1-75 (acceptor/proximal Ub), 0.5 μM UBE1, 15 μM UBE2R1, 10 mM ATP, 50 mM Tris-HCl (pH 7.5), 10 mM MgCl2, and 0.6 mM DTT. The reaction produced K48-Ub3 [Cys-Ub_prox_] and smaller quantity of K48-Ub5 [Cys-Ubprox]. Chains were purified as described previously (Kristariyanto *et al*, 2015).

###### Assembly of K48-Ub_n_ containing Cys residue at the proximal Ub (K48-Ub_n_ [Cys-Ub_dist_])

Chains were generated as described previously (Abdul Rehman *et al*, 2021). Briefly, K48-Ub_n_ [Cys-Ub_dist_] was assembled by generating a ladder of polyUb chains capped with 6His-Ub (K48R, K63C). The assembly reaction was carried out for 18 h at 30°C in a 1 mL volume containing 2500 mM WT ubiquitin, 250 mM (N-term 6His)-Ub (K48R, K63C), 0.5 mM UBE1, 15 mM UBE2R1, 10 mM ATP, 50 mM Tris-HCl (pH 7.5), 10 mM MgCl2, and 0.6 mM DTT. To select for chains that had incorporated the Ub capping mutant, the reaction mix was passed over HisTrap FF 5 mL column (Cytiva) and then fractionated by size over Superdex 16/60 (Cytiva). Fractions containing chains between Ub7-Ub20 were pooled, concentrated and buffer exchanged into PBS.

###### Coupling Cys-containing Ub chains to fluorescent label

An infrared dye, IRDye 800CW Maleimide (LICOR), was conjugated to polyUb chains containing a Cys residue through thiol-maleimide click chemistry. Cys-containing K48-linked polyUb chains of defined length were diluted in 20 mM Tris pH 7.5 and 500 μM TCEP to a final concentration of 50-100 μM. These were purged with argon and incubated at RT for 3 hr. IRDye 800CW Maleimide was dissolved in DMSO to prepare a 5 mM stock. A four-fold excess of fluorescent dye was added to the polyUb chains. The reaction was purged with argon and incubated at RT for 2 h in the dark. The addition of an excess of β-mercaptoethanol quenched the reaction. Labelled polyUb chains were separated from unreacted dye using either PD-10 or Hiprep 26/10 desalting columns (Cytiva) and passed over either Superdex75 16/60 or Superdex200 16/60 (Cytiva).

##### Analysis of recombinant proteins

###### SDS-PAGE

Protein samples were prepared in 4x LDS buffer to a final concentration of 1x and supplemented with 5 mM DTT. Samples were heated to 95°C for 5 mins unless they contained polyUb, which smear when resolved on a gel after such a treatment. Samples were resolved along with Precision Plus 10-250 kDa markers (Bio-Rad) on either NuPAGE 4-12% Bis-Tris gradient gels (Invitrogen) or 15% Bis-Tris gels. Proteins were separated by electrophoresis at 200 V in 1x MES-SDS buffer (ForMedium) for resolution of smaller proteins and 1x MOPS-SDS (ForMedium) buffer for resolution of larger proteins. Gels were then either Coomassie-stained (Instant Blue, Expedeon) or silver-stained (Silver Stain Kit, Pierce).

###### Qualitative deubiquitylation assays

2x stocks of both DUB and substrate were prepared in DUB assay buffer. The DUB was incubated at room temperature for 10 minutes to fully reduce the catalytic cysteine. An appropriate volume of 2x DUB was added to an equal volume of 2x substrate. The reaction mix was incubated at 30°C for the indicated time. 10 μL samples were taken at indicated time points and the reaction was quenched with 4 μL 4x LDS sample buffer. Samples were then resolved by SDS-PAGE and the gels were either silver-stained or Coomassie-stained.

###### Quantitative deubiquitylation assays

For assays using fluorescently labelled Ub chains, 2x stocks of DUB and fluorescently labelled chains were prepared in reaction buffer (50 mM NaCl, 50 mM Tris-HCl pH 7.5, 0.25 mg/mL BME, and 0.25 mg/ml BSA). For the assays, an equal volume of DUB was added to fluorescently labelled chains to final concentrations of 1 μM and 500 nM, respectively. Reactions were carried out at 30°C. 2.5 μL of the reaction mix was added to 1x LDS dye with reducing agent at the indicated time points. Samples were resolved on a NuPAGE 4-12% Bis-Tris gel (ThermoFisher) and visualised on an Odyssey CLx (LI-COR) using the 800 nm channel. Intensities were quantified using Image Studio Lite (LI-COR) and processed using Prism (GraphPad).

###### Pulldown of GST-RAD23A and MINDY3

Equal volumes of the tagged species (7.5 μM) and the untagged species (15 μM) were mixed and incubated on ice for 1h with occasional agitation. GSH resin was added, and the mixture was incubated for a further hour on ice with occasional agitation. This was then added to a polypropylene column and the flow through collected. The resin was washed with 3 x bed volumes of GST_PD buffer. Protein was then eluted from the resin using 3 x bed volumes of GST_PD buffer supplemented with 10 mM reduced glutathione. Samples of each fraction were then mixed with 4x LDS dye and then resolved by SDS-PAGE. Gels were Coomassie stained.

###### Analytical gel filtration for complex analysis

A 60 μL mix was prepared containing 10 μM of each species in SEC_2 buffer. This was incubated for 1hr on ice before being injected onto a Superdex75 3.2/300, Superdex 200 3.2/300, or Superose 6 3.2/300. Fractions were collected and analysed by SDS-PAGE with Coomassie staining.

###### Isothermal Titration Calorimetry

ITC measurements were performed on MicroCal PEAQ-ITC (Malvern) at 25°C. Prior to measurements, all proteins were passed over Superdex 75 16/60 (Cytiva) and then dialysed into a buffer containing 50mM Tris-HCl pH 7.5, 150mM NaCl and 250 µM TCEP. The DUB was taken up by the syringe and was titrated into the cell which contained K48-linked polyUb chains (K48-Ub2, Ub3 or Ub4 or Ub5). Likewise, in other experiments, the scarcest species was added into the cell to conserve resources. 2 µl of injectant was dispensed in 4-sec duration with 130-sec spacing between injections for a total of 16 injections. Data were analysed and titration curves were fitted using MicroCal PEAQ-ITC (Malvern) analysis software (n = 2; mean ± SD).

###### Pulldown of recombinant ubiquitin chains and UBL domains with HALO-tagged EF-hand

A HALO-tagged MINDY3^EF-hand^ fusion was used for pulldowns with either recombinant ubiquitin chains or isolated UBL domains. Briefly, 10 nmol of HALO-EF-hand was immobilised on 100 µl of HALOlink resin (Promega) in coupling buffer (50 mM Tris-HCl pH 7.5, 150 mM NaCl, 0.5% NP-40, 0.5 mM TCEP) at 4°C overnight. Beads were washed thrice with coupling buffer and resuspended in 100 µl of pulldown buffer (50 mM Tris-HCl pH 7.5, 150 mM NaCl, 0.1% NP-40, 0.5 mM TCEP, 0.5 mg/ml BSA). Per reaction, 20 µl of bead slurry was mixed with 30 pmol of prey protein in 480 µl of pulldown buffer and samples were incubated on a rotator at 4°C for 2 hours. Beads were washed thrice with wash buffer (50 mM Tris-HCl pH 7.5, 150 mM NaCl, 0.2% NP-40, 0.5 mM TCEP) and eluted with LDS sample buffer (ThermoFisher). Samples were subsequently analysed by SDS-PAGE and silver staining (ThermoFisher).

### Protein Crystallisation and Data Collection

#### MINDY3apo

Initial MINDY-3 crystals were obtained using sitting-drop vapor diffusion method in a 96-well format. Subsequently, crystals were reproduced and refined in a hanging-drop vapor diffusion setup. Drops containing 15.8 mg/mL MINDY-3 FL were mixed 1:1 with mother liquor composed of 20% (v/v) PEG 6000 and MES pH 6.0. Crystals appeared after two days. Crystals were cryoprotected using 25% ethylene glycol and snap-frozen in liquid nitrogen. SeMet-labelled MINDY3 was crystallized under identical conditions. A native dataset was collected at beamline MX-14.2 at the BESSY II synchrotron (Berlin, Germany). A dataset for experimental phase determination by SAD was acquired at beamline P11 at PETRA III (Hamburg, Germany).

#### MINDY3^EF-hand^ + RAD23A^UBL^

MINDY3 (297-370) and RAD23A (1-84) were separately expressed in Rosetta2(DE3) cells and purified via their GST tags, and subsequently by SEC. The two species were mixed in a 1:1 ratio giving a final concentration of 15 mg/mL. Crystals grew in sitting drops containing a 1:1 ratio of protein to mother liquor which contained 0.2 M potassium chloride and 2.2 M ammonium sulphate. Crystals were frozen in cryoprotectant consisting of mother liquor supplemented with 35% glycerol. Diffraction data were collected at I04 beamline, DLS, UK.

### Structure Determination and Refinement

#### MINDY3apo

Initial attempts to solve MINDY-3 structure by molecular replacement (MR) using structurally characterized homolog MINDY-1 (or subdomains) were unsuccessful. Sequence independent MR tool SIMBAD using MoRDa database also could not solve the phase problem, advocating for single-wavelength anomalous diffraction (SAD). Native and anomalous datasets were processed with the XDS software package (Kabsch, 2010). The structure of SeMet-labelled MINDY-3 was determined using AutoSol at 2.95 Å, which identified strong selenium sites for 9 out of 11 methionines, which were used as anchors for initial model building. The resulting model was converted to poly-alanine to minimize side chain bias and used as a template for MR in Phaser-MR, which solved native MINDY-3 structure at 2.3 Å resolution. Refinement was performed using phenix.refine (Afonine *et al*, 2012) with iterative rounds of model building in Coot (Emsley *et al*, 2010).

#### MINDY3^EF-hand^ + RAD23A^UBL^

Diffraction images were processed with the autoPROC pipeline (Vonrhein *et al*, 2011) using XDS (Kabsch, 2010), Pointless (Evans, 2006) and Aimless (Evans & Murshudov, 2013). The structure was solved by molecular replacement (Phaser) (McCoy *et al*, 2007)) using AlphaFold predicted models of each entity as search models. The asymmetric unit contained two copies of the EFH and two copies of the UBL, but only one copy of the complex. The model was refined with phenix.refine of the Phenix software suite (Afonine *et al*, 2012) and through manual building in COOT (Emsley *et al*, 2010).

##### AlphaFold predictions

AlphaFold models were generated by inputting protein sequences into the Colab AlphaFold notebook platform (Jumper *et al*, 2021)

##### Data analyses and figure generation

Protein sequences were acquired from Uniprot. DNA sequence analyses were carried out in Snapgene. Protein amino acid sequences were obtained from Uniprot (https://www.uniprot.org). Sequence alignments were generated using the Clustal Omega multiple sequence alignment program in Jalview (Waterhouse *et al*, 2009). SDS-PAGE gels using fluorescently labelled species were imaged using a LICOR Odyssey CLx and analysed in ImageStudioLite. Graphs were prepared in GraphPad Prism (https://www.graphpad.com). Structure representations were generated in PyMol (https://pymol.org/2/) and ChimeraX (Pettersen *et al*, 2021). All figures were generated and annotated in Adobe Illustrator.

##### Mammalian cell culture

U2OS Flp-In Trex and HEK293 Flp-In Trex cell lines were maintained in DMEM (Gibco) and RPE-1 cells were maintained in DMEM/F12 1:1 (Gibco). Both media types were supplemented with 10% FBS (Gibco), 2 mM L-glutamine (Gibco), and 100 U/ml Penicillin/Streptomycin (Gibco), and cells were incubated at 37°C with 5% CO2 unless otherwise stated. Trypsin(0.05%)-EDTA (Gibco) was used to dissociate cells for passage. All cell lines were routinely tested for mycoplasma.

Cell lines were transiently transfected with vectors using PEI Max 40k (Polysciences) at a 1:4 ratio (w/w) of DNA:PEI. Following transfection, cells were incubated at 37°C for 48 hours prior to harvest.

For generation of cell lines stably expressing tetracycline-inducible GFP-tagged constructs, Flp-In Trex cells were co-transfected with a 1:9 ratio (w/w) of GFP vector:pOG44 Flp recombinase vector using Lipofectamine LTX (ThermoFisher) according to the manufacturer’s protocol. To select for integrant cells, 24 hours post-transfection, media was switched out for fresh DMEM supplemented with 200 μg/ml hygromycin B. Selection media was periodically refreshed, and cultures monitored until all mock transfected control cells were dead. Tet-inducible expression of the proteins of interest were subsequently confirmed via western blotting with an anti-GFP antibody, following overnight incubation with 1 μg/ml tetracycline.

The RPE-1 and U2OS MINDY3 KO cell lines were generated by co-transfection of paired CRISPR guides and dCas9 nickase.

### RNA interference

RNAi was carried out using Lipofectamine RNAiMAX (Thermo Scientific) according to the manufacturer’s protocol. Briefly, cells were seeded into 6-well plates at 1-2×10^5^cells/well. The following day, cells were transfected with 25 pmol of siRNA duplexes prepared in RNAiMAX reagent. Cells were then incubated at 37 °C for 48 hours prior to harvest and subsequent analysis. All siRNAs were sourced from Dharmacon and sequences are presented in **Table 4**.

### Crosslinking immunoprecipitation

Following harvest, cells were treated with 1 mM DSP (ThermoFisher) for 30 minutes at room temperature then quenched with 50 mM Tris-HCl, pH 7.5 and incubated for a further 20 minutes. Cells were pelleted at 1000 x g for 10 minutes, resuspended in 1% SDS lysis buffer (1% SDS in 1XTBS, 1X cOmplete PIC, 0.02% benzonase) and incubated at 37°C for 15 minutes. SDS concentration was diluted to 0.2% with Co-IP lysis buffer and samples clarified via centrifugation. Resulting supernatants were mixed with 20 μl of FLAG affinity gel (Merck) and incubated on a rotator at 4°C for 1 hour. Beads were washed four times with Co-IP lysis buffer and eluted in four bead volumes of 2XLDS sample buffer lacking reducing agent. Elution fractions were separated from beads by applying to SpinX filter columns and spun at 2,500 x g for 2 minutes. Input and elution fractions (± 50 mM DTT) were subsequently analysed by SDS-PAGE followed by immunoblotting.

### Western blotting

Protein samples were mixed with 4X LDS Sample buffer and 10X Reducing Agent (both Thermo Scientific) and incubated at 70 °C for 10 minutes. Following SDS-PAGE and protein transfer, membranes were stained with Ponceau S (Sigma) to assess loading and transfer efficiency. Chemiluminescent blots were subsequently visualized via a ChemiDoc MP (BioRad) using Clarity or ClarityMAX ECL reagents (BioRad), and fluorescent blots with an Odyssey CLx (LiCor Biosciences).

### Antibodies

Antibodies were sourced from the indicated manufacturers and used at 1:2000 dilution unless otherwise stated: anti-FLAG HRP (Sigma, A8592, 1:10000), anti-DYKDDDDK HRP (ProteinTech, HRP-66008, 1:10000), anti-GFP (ProteinTech, 50430-2-AP, 1:10000), anti-RAD23A (CST, 24555), anti-RAD23B (ProteinTech, 67988-1-Ig), anti-MINDY3 (Eurogentec, R179), anti-MINDY3 (MRCPPU Reagents & Services, DA026 2^nd^ bleed), anti-Ubiquitin (Biolegend, 646302), anti-GAPDH (CST, 2118, 1:5000), anti-CPD (abcam, Ab10347), anti-cDNA (abcam, Ab27156). Secondary detection was carried out using anti-rabbit (CST, 7074), anti-mouse (CST, 7076) or anti-sheep (R&D Systems, HAF016) HRP-conjugated antibodies (all three 1:5000), or anti-rabbit, anti-mouse or anti-goat IRDye800CW/680RD-conjugated (LiCor Biosciences, 926-32211, 926-32210, 926-32214, 926-68073, 926-68070, 926-68074, all 1:15000) antibodies.

### UV-laser striping

Laser micro-irradiation was performed using a Leica Stellaris 8 confocal microscope (Leica microsystems) equipped with a white light laser and an environmental chamber set at 37°C, supplemented with 5% CO2. Image acquisition and irradiation was performed using a sequential acquisition pipeline built using the ‘Live Data Mode’ module within the Leica LASX acquisition software (v4.8.1.29271). Software-based autofocus was first performed to locate the cells of interest. A pre-irradiation image was then recorded, followed by 405nm laser micro-irradiation. Irradiation was performed by scanning at low 16×16 frame resolution resulting in a defined pattern of 16 horizontal scan lines across the field of view. The 405nm laser was set to 100% power and scanned bi-directionally across the sample at 3Hz and iterated by using 32x averaging. A time-lapse was subsequently performed, acquiring images every 30 seconds for 10 minutes to follow the dynamics of the protein of interest after irradiation. For pre- and post-irradiation image acquisition, 16-bit images were acquired bi-directionally at a frame size of 1024×1024 with a scan speed of 600Hz, with 2x line averaging and the pinhole set to 1AU. GFP was excited using the white light laser at 477nm. Z-stacks were acquired with 6 1-um optical sections per frame. For collection, a HyD X detector was used in digital mode, with the minimum gain setting (10) and images were acquired using 2x accumulations per frame, collecting between 483-814nm. Experiments were performed using a Leica HC PL APO CS2 63x/1.40 oil immersion objective with Leica’s Type F immersion oil.

### Yeast-2-Hybrid Screening

Y2H screening was performed by Hybrigenics Services (France) using full-length MINDY3 (purified and with GST tag cleaved as described above) as a bait and a human placenta cDNA library as prey.

## Data Availability

The protein crystal structures generated in the study have been deposited as follows:

- MINDY3 apo structure: PDB 9RMD
- MINDY3^EF-hand^:RAD23A^UBL^ structure: PDB 8Q06

## Acknowledgements

We thank members of the Kulathu lab for helpful discussions. We thank Profs. Dantuma (Karolinska) and Rouse (Dundee) for helpful discussions. We thank Drs Lange and Magnussen for help with AlphaFold predictions, Dr Gregorczyk for assistance with UV-laser stripe live cell imaging and image processing, Dr Montava Garriga for the U2OS MINDY3 KO cell line, and Dr Shen, Dr Knebel and Ms. Johnson for protein reagents. This work was supported by funding from MRC grant MC_UU_00018/3 and MC_UU_00038/3 (YK), ERC Consolidator grant (grant 101002428; YK), Marie Skłodowska-Curie Innovative Training Networks (ITN) UbiCODE (765445) and by the OPUS24 grant (2022/47/B/NZ1/01941; S.G.) from the National Science Centre (NCN, Poland).

## Author contributions

### Credit role taxonomy

Conceptualization – SG, YK

Data curation

Formal analysis

Funding acquisition – SG, YK

Investigation – LAA, MRM, RK, ROD, MG, TC

Methodology

Project administration

Resources

Software

Supervision – SG, YK

Validation

Visualization – LAA, MRM, ROD

Writing – original draft – LAA, MRM, YK

Writing – review & editing – LAA, MRM, SG, ROD, YK

## Supplementary Figures

**Figure EV1:**
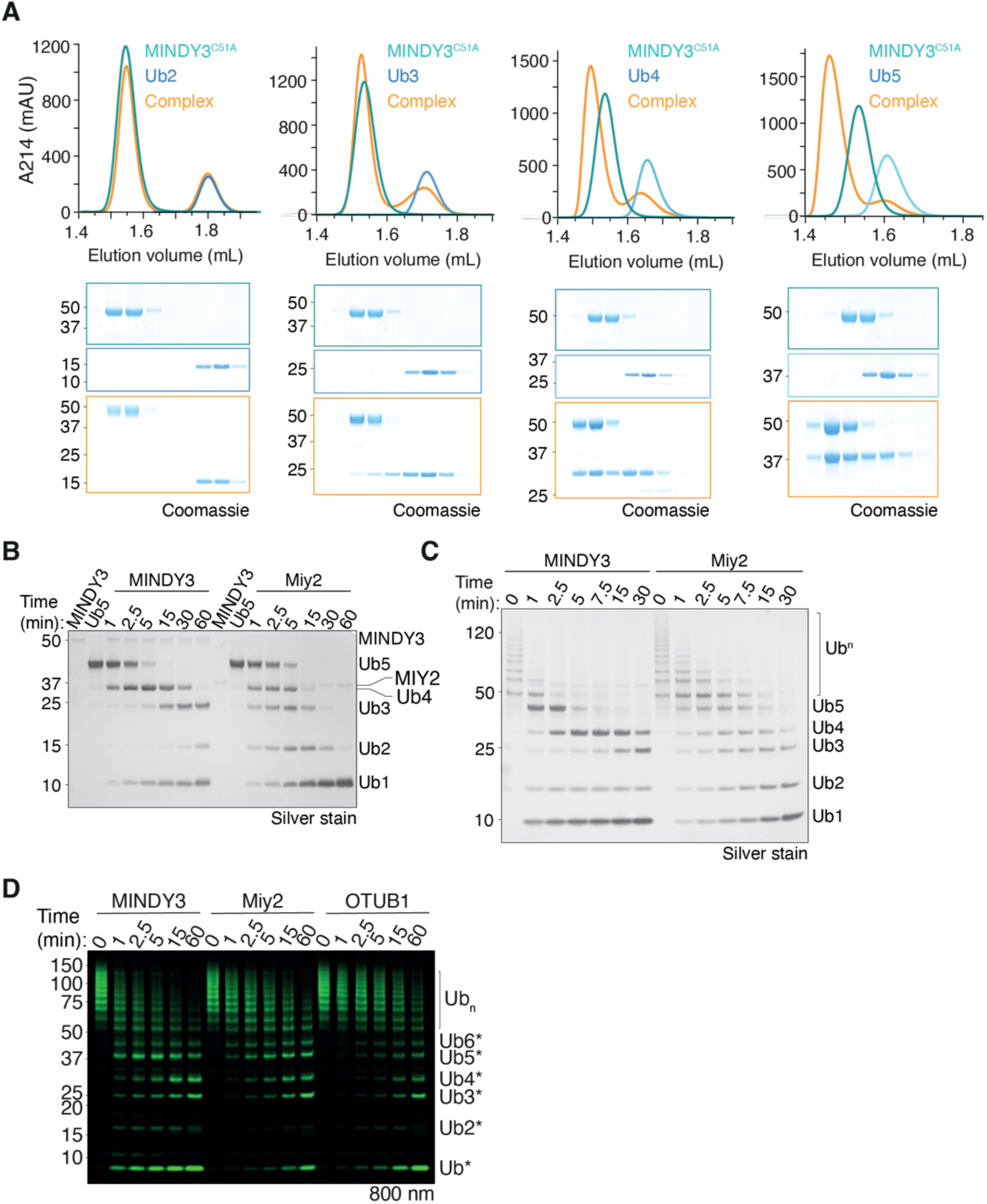
**A)** Analytical SEC of MINDY3^C51A^ with K48-linked Ub2-5 after pre-incubation. **B)** DUB assay of MINDY3 against K48-linked Ub6. **C)** DUB assay of MINDY3 against K48-linked polyUb of length >Ub6. **D)** DUB assay of MINDY3 against K48-linked polyUb of length Ub6+ labelled at the extreme distal moiety with a fluorophore. Miy2 and OTUB1 are endo-DUB controls.

**Figure EV2:**
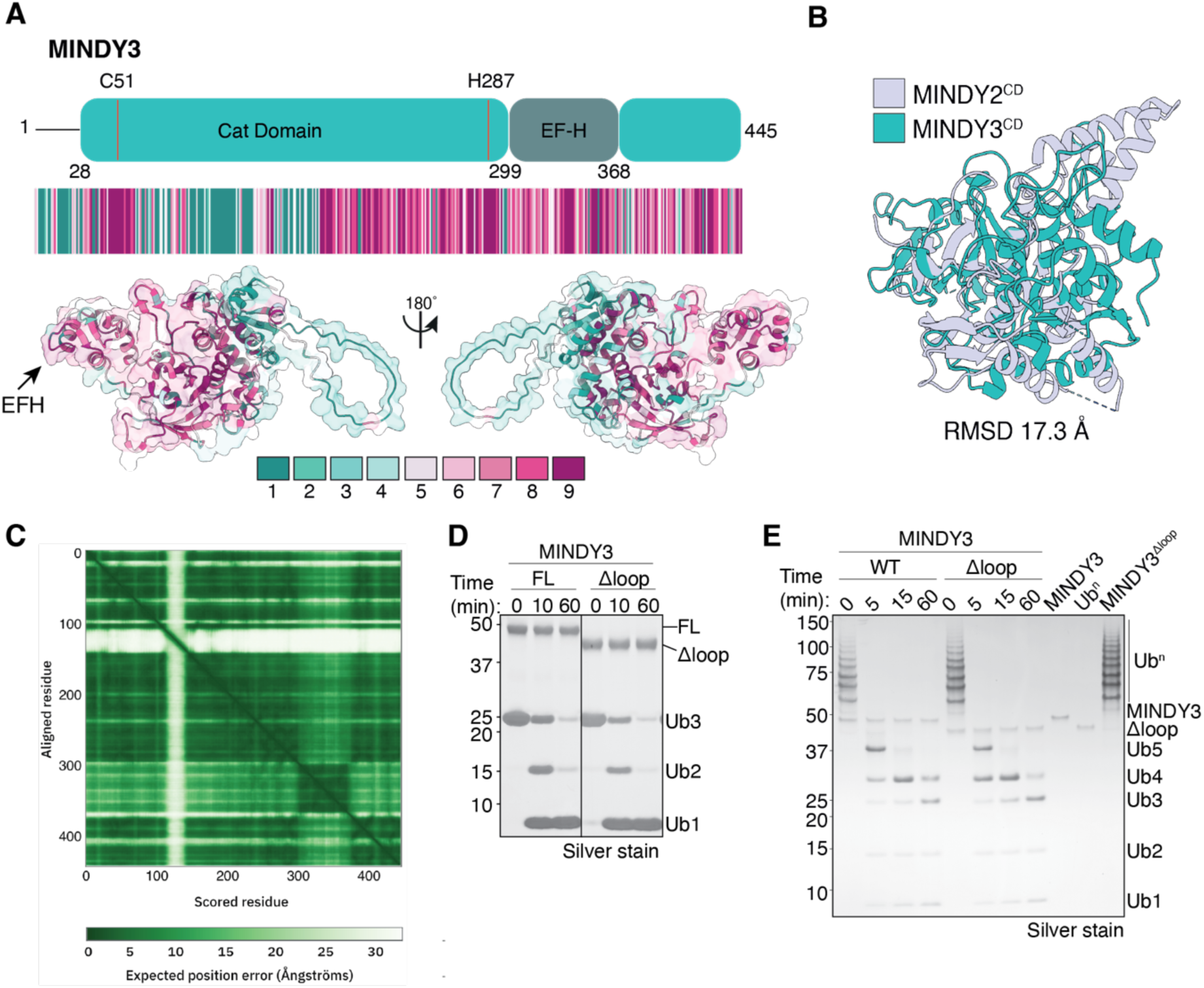
**A)** Conservation analysis of each amino acid position of MINDY3 generated by Consurf (Ashkenazy *et al*, 2016) from the sequences of 186 vertebrate species. Each residue is given a score between 1-9, with 1 being the least conserved and 9 being the most conserved. Consurf analysis is then mapped onto the AlphaFold prediction for FL MINDY3. **B)** Superposition of the crystal structure of MINDY3 with the crystal structure of the catalytic domain of MINDY2 (PDB: 6Z49). **C)** PAE plot of the AlphaFold prediction of full length MINDY3. Low confidence regions correlate to the flexible loop region in MINDY3. **D)** DUB assay of MINDY3 and MINDY3^Δloop^ against K48-linked Ub3. **E)** DUB assay of MINDY3 and MINDY3^Δloop^ against K48-linked polyUb of Ub6+.

**Figure EV3:**
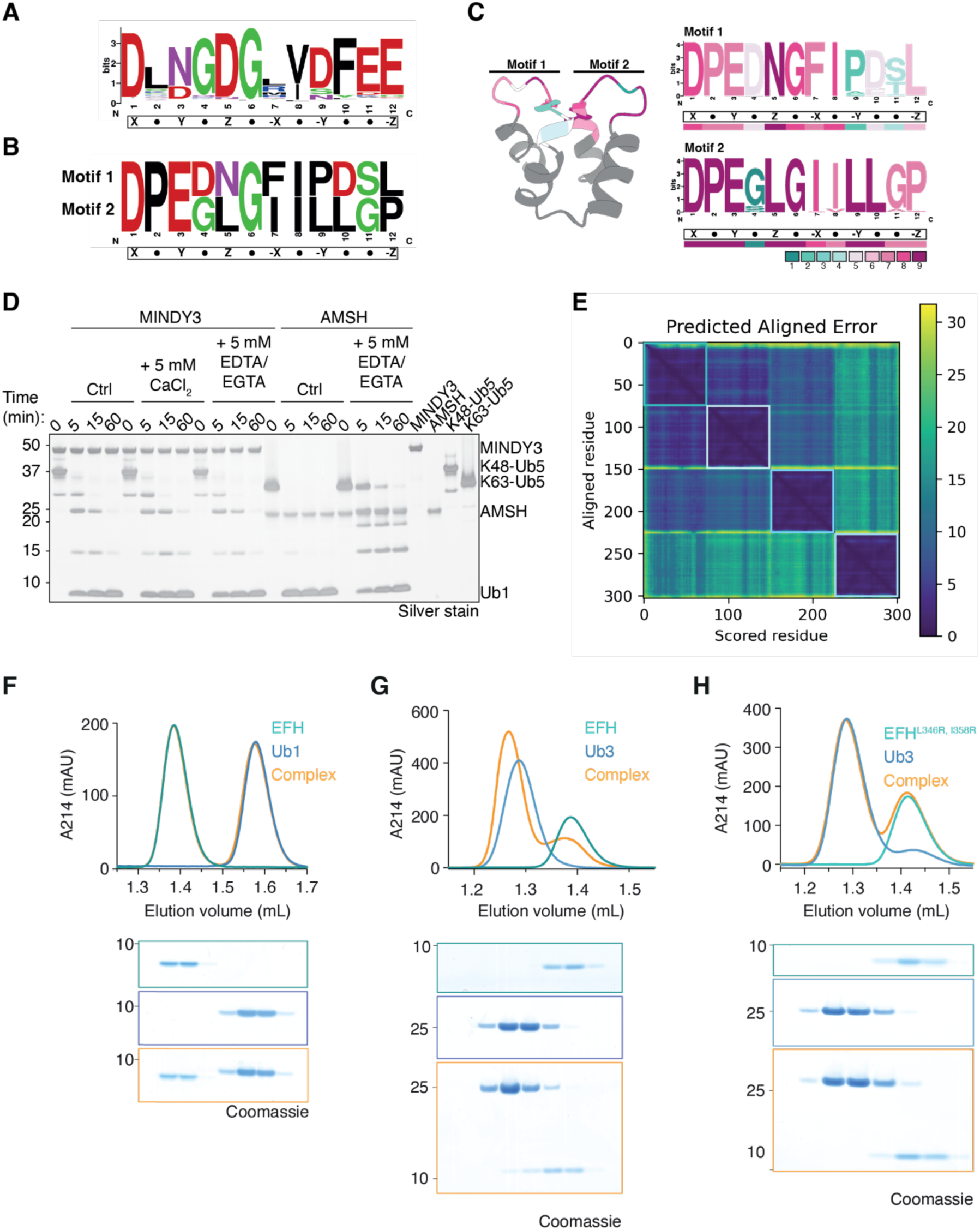
**A)** WebLogo visualisation of sequence conservation of the EF-hand motif generated from 878 vertebrate genes. The overall height of a letter stack indicates the level of conservation at that position, while the height of an individual letter indicates the relative frequency of the amino acid at that position (Crooks *et al*, 2004). **B)** WebLogo visualisation comparing the two EF-hand motifs in MINDY3. The height of an individual letter indicates the relative frequency of the amino acid at the position. **C)** Left: MINDY3^EF-hand^ with the EF-hand motifs coloured according to Consurf analysis of MINDY3 from 186 vertebrate genes. Right: As in (A) but with MINDY3 from 186 vertebrate genes. Coloured by Consurf analysis. **D)** DUB assay in which MINDY3 was untreated or pre-incubated with CaCl_2_ or EDTA/EGTA prior to incubation with K48-linked Ub5. AMSH, a metalloprotease, was used as a control with K63-linked Ub5. **E)** PAE plot for the AlphaFold model of MINDY3^EF-hand^ + three ubiquitin molecules. Coloured boxes indicate individual proteins as in figure 3D. **F-H)** SEC of MINDY3^EF-hand^ with monoub (F) or K48-Ub3 (G), or a mutant MINDY3^EF-hand^ with K48-Ub3 (H).

**Figure EV4:**
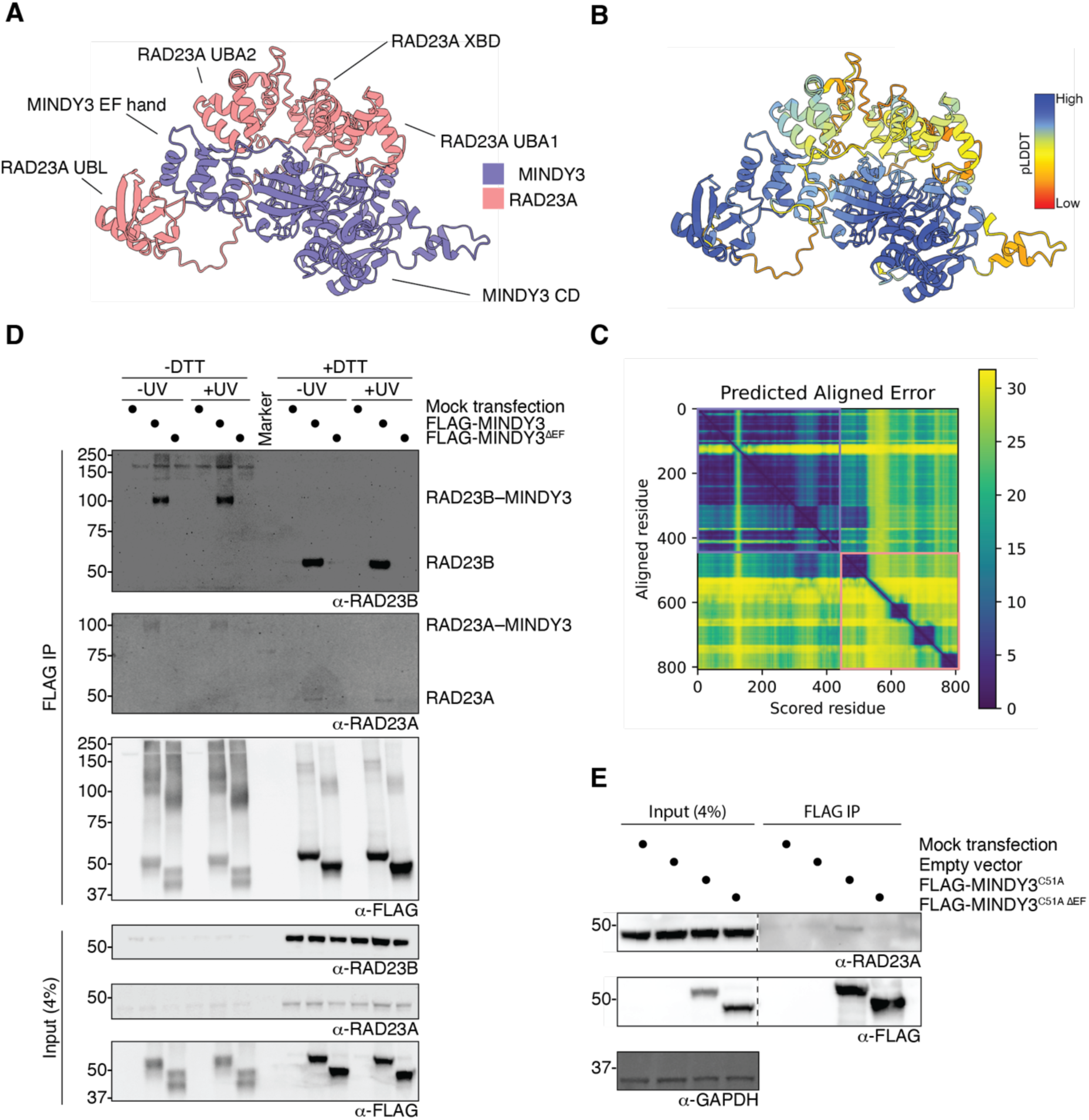
**A)** AlphaFold 2 prediction of full-length MINDY3 with full-length RAD23A. **(B)** Alphafold 2 prediction as in (A), but coloured according to confidence based on pLDDT score**. (C)** Associated PAE plot for AlphaFold prediction. Coloured boxes indicate individual proteins as in figure EV4A. **(D)** Crosslinking IP of MINDY3 from HEK293 cells. Cells were transiently transfected with FLAG-MINDY3 ± EF-hand and crosslinked with DSP before FLAG IP. Input and elution fractions (± DTT to resolve crosslinks) were analysed via western blotting with indicated antibodies. **(E)** Co-immunoprecipitation of MINDY3^C51A^ and RAD23A from HEK293 cells. Cells were transiently transfected with catalytically dead FLAG-MINDY3 ± EF-hand before FLAG IP. Input and elution fractions were analysed via western blotting with indicated antibodies.

**Figure EV5:**
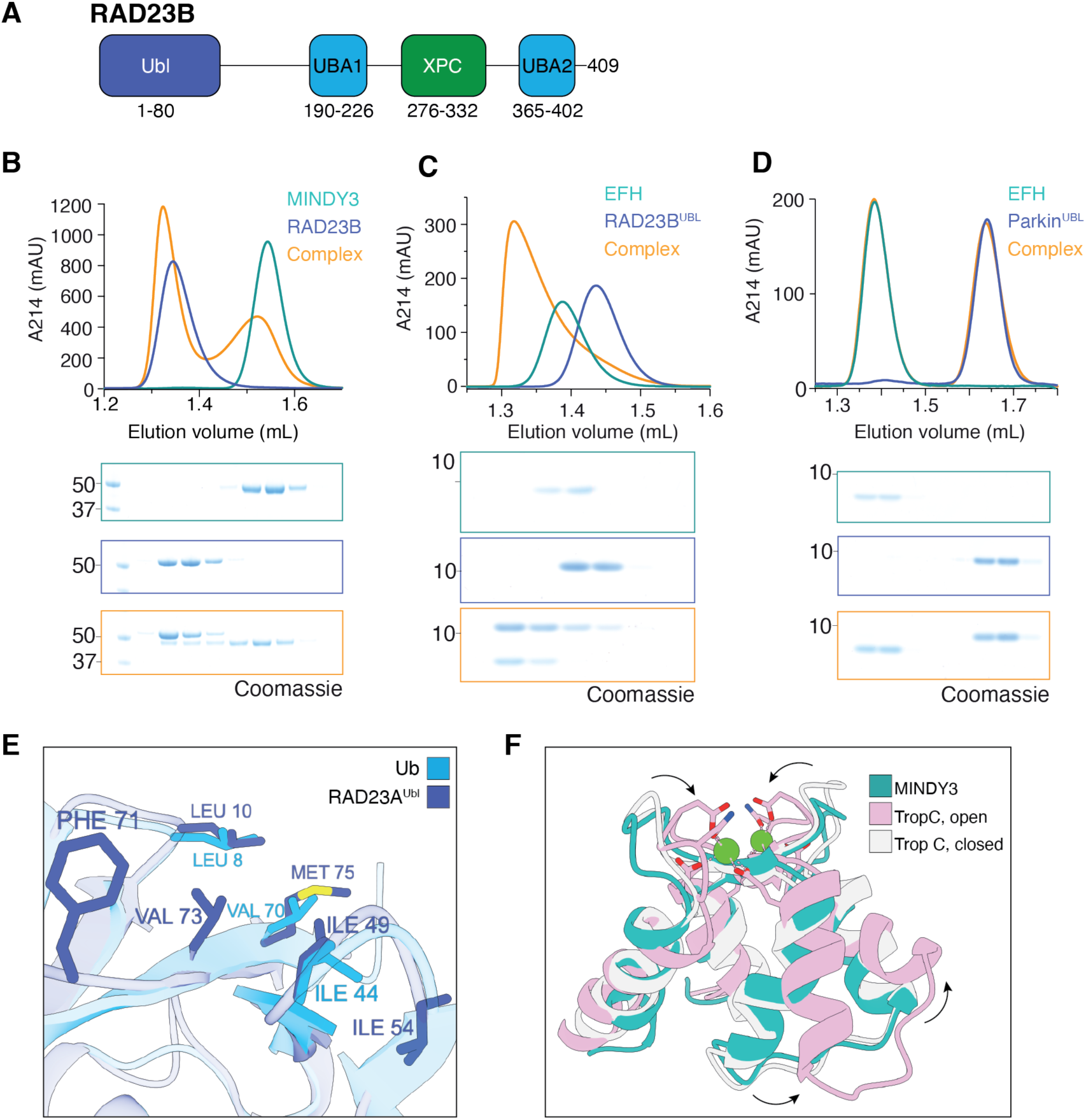
**A)** Domain schematic of RAD23B showing the N-terminal UBL domain, the two UBA domains and the XPC-binding domain. **B)** SEC analysis of MINDY3 and FL RAD23B. **C)** SEC analysis of MINDY3^EF-hand^ with RAD23B^UBL^. **D)** SEC analysis of MINDY3^EF-hand^ with Parkin^UBL^. **E)** Comparison of the interaction interface of RAD23A^UBL^ with the I44 patch of Ub. **F)** Transition between open and closed forms of the EF-hand in Troponin C (PDB: 5TNC) superposed with MINDY3^EF-hand^.

**Figure EV6:**
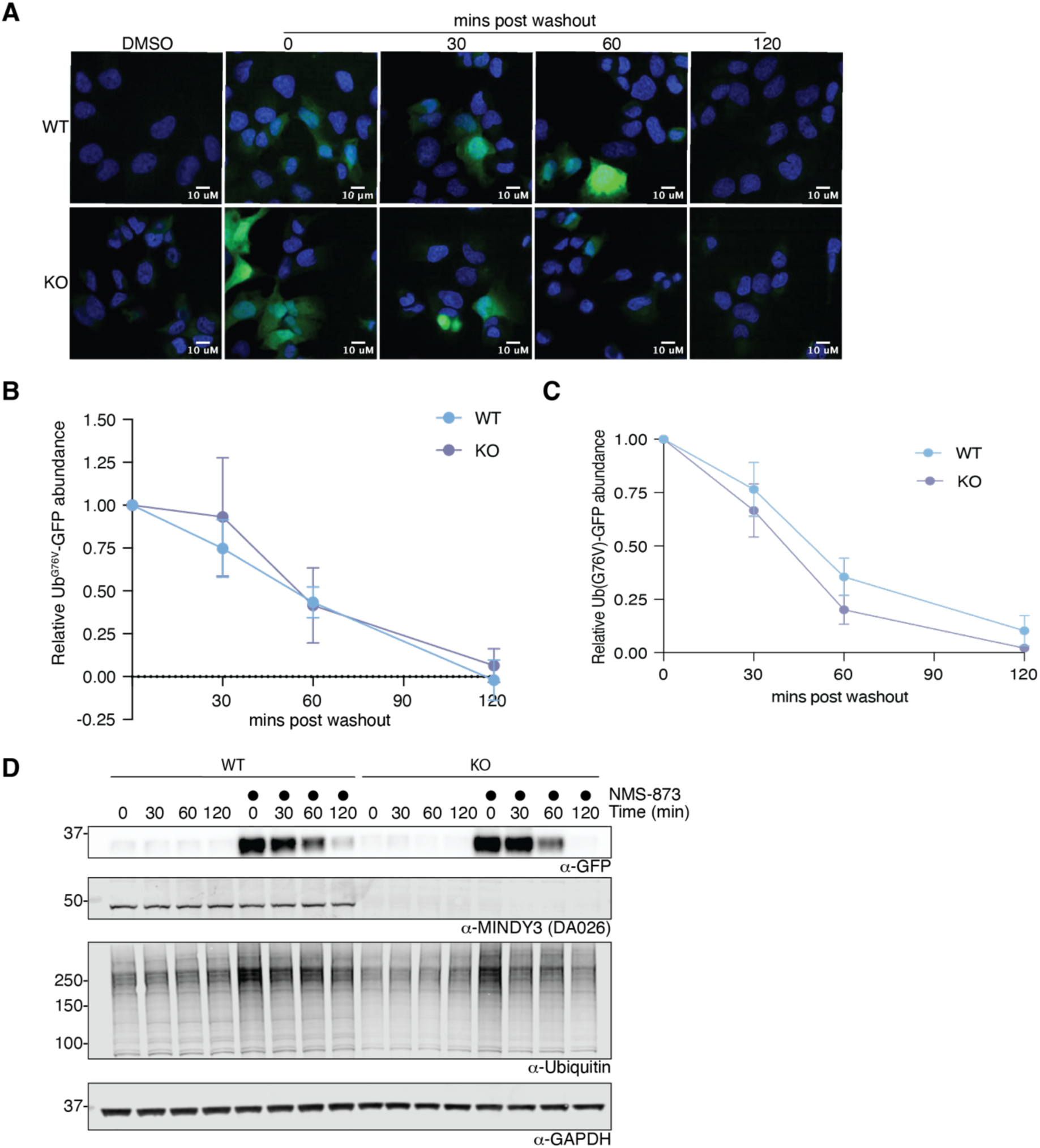
**(A)** Representative images of Ub^G76V^-GFP abundance in WT or MINDY3 depleted U2OS cells. To prevent the degradation of Ub^G76V^-GFP cells were treated with the p97 inhibitor NMS-873 for 2 h. Degradation of Ub^G76V^-GFP was assessed by fluorescence microscopy at the indicated times after NMS-873 washout. **(B)** Quantification of the GFP signal detected in (A). Data points shown are mean nuclear GFP intensities normalised to 0% and 100% corresponding to DMSO and NMS treated samples respectively. Three independent experiments, with a minimum of 500 cells per condition were quantified. **(C)** Relative stability of Ub^G76V^-GFP in WT and MINDY3 KO U2OS cells. Cells were pre-treated with NMS-873 for 4 hours, and Ub^G76V^-GFP abundance was then assessed at the indicated time by western blot and normalised for loading with corresponding GAPDH levels. Error bars indicate SD based on three independent experiments each with a minimum of two replicates. **(D)** Representative western blot of data presented in (C).

**Figure EV7:**
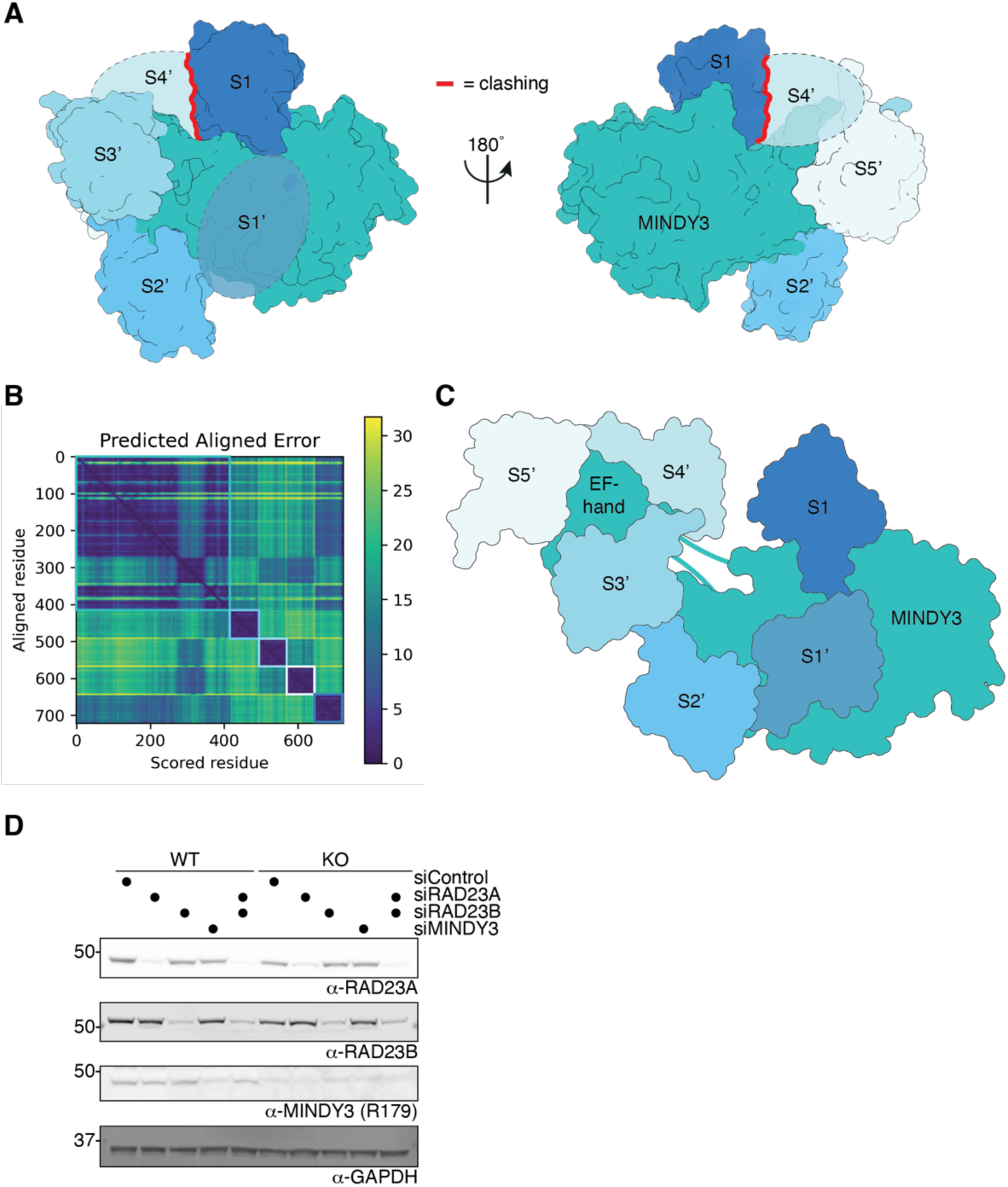
**A)** AlphaFold model of MINDY3 with 4 ubiquitin molecules where Ub is placed at the S1, S2’, S3’ and S5’ sites. A ubiquitin can be placed at the putative S1’ site based on geometry and the distance between the S1 and S2’ ubiquitin molecules. The S3’ and S5’ ubiquitin molecules correspond to the distal and proximal ubiquitins in the EF-hand:UBL model in Fig. 3D. Modelling the medial ubiquitin onto this AlphaFold model results in extensive clashing with the S1 ubiquitin. **B)** PAE plot for the AlphaFold model of MINDY3 with four ubiquitin molecules. Coloured boxes indicate individual proteins as in figure EV7C. **C)** Schematic of MINDY3 bound to 6 ubiquitin molecules. The core catalytic domain binds to 3 ubiquitins, whilst the EF-hand binds the remaining 3. The steric clash between the S4’ and S1 ubiquitins seen in **(Fig. EV7A)** is avoided by repositioning the EF-hand via the flexible linkers that connect it to the core catalytic domain. **D)** MINDY3 and RAD23A/B do not affect each other’s abundance. WT and MINDY3 KO RPE-1 cells were transfected with siRNAs targeting MINDY3, RAD23A or RAD23B, and protein levels assessed via western blot using indicated antibodies.

